# Age-related difference in gray matter volumes is heterogeneous across cerebellar regions but neither related to the topological organization nor the associated sensorimotor function

**DOI:** 10.64898/2026.02.13.705695

**Authors:** Anda de Witte, Anouck Matthijs, Caroline Nettekoven, Jolien Gooijers, Jean-Jacques Orban de Xivry

**Author notes:** **Corresponding author** Jean-Jacques Orban de Xivry, Tervuursevest 101, 3001 Leuven, Belgium.

## Abstract

Aging affects cerebellar structure, yet the regional specificity of this decline and its relationship to sensorimotor functions remain unclear. In this study, we quantified age-related cerebellar gray matter differences in 50 young and 80 older adults using both anatomically defined cerebellar parcellations and a functionally defined cerebellar atlas. Across both anatomical and functional parcellations, older adults showed robust reductions in gray matter volume relative to young adults. This age-related gray matter vulnerability was region-specific as age differences varied significantly across regions: both the current dataset and the additionally used Cam-CAN dataset demonstrated a larger negative effect of age in the posterior than the anterior lobe, and a smaller difference in the action domains than in the other functional domains. Structural covariance analyses revealed that correlations between cerebellar regions were determined primarily by spatial proximity and, to a lesser extent, by medial–lateral (vermis–hemisphere) organization or functional similarity. Importantly, the topological organization of the cerebellum did not differ across age groups, indicating preserved structural patterns despite widespread gray matter loss. Finally, despite substantial interindividual variability in behavioral, regional cerebellar gray matter volumes, whether anatomically or functionally defined, did not predict inter-individual variability for any of our eight cerebellum-dependent outcomes. This absence of structure–function relationship suggests that behavioral performance is maintained through compensatory mechanisms or microstructural features not captured by regional gray matter volume. Together, the results suggest heterogeneous age-related cerebellar degeneration alongside preserved topological organization and no measurable impact on cerebellar sensorimotor function, supporting the notion of a robust cerebellar reserve throughout healthy aging.

## 1. Introduction

Although the cerebellum is often called the “little brain”, it contains more than half of all neurons in the human brain (Andersen et al., 1992; Herculano-Houzel, 2009). The cerebellar gray matter, which contains the neuronal cell bodies, is central to many functions, particularly through the activity of Purkinje cells. These cells are vital for motor control and coordination due to their unique capacity to integrate sensory and motor signals during self-generated motion (Zobeiri and Cullen, 2024). Loss of Purkinje cells, as observed in cerebellar ataxia patients with gray matter atrophy, is thought to contribute to disruptions in sensorimotor behavior. These neuropathological changes are consistent with the balance impairments and fine motor deficits commonly observed in affected individuals (Jäschke et al., 2023; Liu et al., 2023).

Although most evident in pathological conditions such as ataxia, cerebellar gray matter loss is not limited to disease, as reductions in cerebellar gray matter volume and neuronal cell count have also been reported in healthy older relative to younger adults (Andersen et al., 2003; Bernard and Seidler, 2013; De Witte et al., 2026a; Hulst et al., 2015). Interestingly, cerebellar volume in older adults was found to correlate positively with grip strength (Koppelmans et al., 2015) and gait speed (Nadkarni et al., 2014). Another study on physical frailty at older age showed that smaller gray matter volume in the cerebellum was associated with motor-related components of frailty, like muscle weakness and movement slowness (Chen et al., 2015). However, other studies have found that reductions in cerebellar gray matter volume in older adults were unrelated to age-related declines in sensorimotor functions such as movement anticipation and adaptation (De Witte et al., 2026a; Duchek et al., 1994; Filip et al., 2019; Matthijs et al., 2025).

To better understand the relationship between cerebellar gray matter and cerebellum-related sensorimotor behavior, it is insufficient to treat the cerebellum as a single, homogeneous structure. Within the cerebellum, three major lobes can be identified: the anterior, posterior, and flocculonodular lobes, each separated by distinct fissures. Additionally, the cerebellum is divided into two hemispheres on either side of the vermis. Beyond these divisions, both vermis and hemispheres can be further portioned into ten lobules. Distinct cerebellar regions participate in different large-scale brain networks and support dissociable functions, including motor coordination, balance, and cognitive processes (Kawabata et al., 2022; O’Reilly et al., 2010). As a result, age-related changes may manifest differently across these regions and may therefore show region-specific relationships with sensorimotor behavior.

It has been widely suggested that the effect of age on cerebellar structure vary across cerebellar subregions (Han et al., 2020). For instance, Andersen et al. (2003), Bernard and Seidler (2013), and Hulst et al. (2015) reported that advancing age primarily affects gray matter volume in the anterior lobules, whereas most regions of the posterior cerebellum showed no specific age-related decline. Another study found similar atrophy patterns in the anterior cerebellum, but also in the vermis and medial sections of the posterior lobe (Wang et al., 2024). Additionally, when subdividing the cerebellum into four transverse zones, it was found that age-related decline was most pronounced in the central zone, encompassing posterior lobules VI and VII, and to a lesser extent in the anterior zone (Ghiyamihoor et al., 2025). This contrast with earlier findings was explained as the study’s use of cytoarchitectural and functional cerebellar subdivisions, which offer greater specificity than traditional anatomical lobular divisions and therefore reveal different patterns of age-related decline. Lastly, (d’Oleire Uquillas et al., 2026) suggested that the impact of aging on anatomical regions forms a gradient, increasing from anterior lobules I-III to posterior lobe crus I, and decreasing from crus I to lobule X. Altogether, these findings suggest that aging may affect cerebellar gray matter unevenly across regions; however, the subtle differences in the literature underscore the need to confirm this pattern. Moreover, it remains unstudied to what extent regional gray matter volume is related to functional performance in the context of aging.

Inter-regional difference in age-related changes in cerebellar structure might be important to understand changes in function with age. Evidence from clinical and brain-stimulation studies provides an important functional framework for interpreting the effects of regional gray matter differences, indicating that gray matter volume in the anterior cerebellar lobe is primarily associated with motor functions (Kansal et al., 2017; Schmahmann, 2012), whereas the posterior lobe is more strongly linked to cognitive and emotional processing (Ciricugno et al., 2024; Stoodley and Schmahmann, 2010). In the healthy aging literature, little attention has been given to how gray matter volume in specific cerebellar regions relates to sensorimotor and cognitive function. Koppelmans et al. (2017) reported that gray matter volume in the cerebellar anterior lobe and the superior portion of the posterior lobe was positively associated with motor skill performance, whereas gray matter volume in the inferior portion of the superior posterior lobe and the inferior posterior lobe was positively associated with cognitive performance, thereby supporting associations previously reported in clinical studies (Ciricugno et al., 2024; Kansal et al., 2017; Schmahmann, 2012; Stoodley and Schmahmann, 2010). Similarly, Bernard and Seidler (2013) reported that working memory and reaction time were positively correlated with gray matter volume in the posterior cerebellum, whereas motor timing was primarily associated with the volume of the vermis of older adults. However, some of their findings diverged from structure-behavior associations found in earlier patient studies. They found that larger gray matter volume in the posterior cerebellum and Crus I was associated with poorer balance, an unexpected result, given that sensorimotor performance is typically linked to the anterior cerebellum rather than posterior regions, and that larger regional volumes are generally assumed to support better function (Stoodley and Schmahmann, 2010). A similar contrasting finding, that motor performance was associated with posterior cerebellar lobe volume in older adults, was also reported by Nadkarni et al. (2014). However, this association disappeared after controlling for information-processing ability, suggesting that motor performance in older adults may partly reflect underlying cognitive components. Altogether, these findings indicate that the functional role of cerebellar regions in motor and cognitive performance is not fully resolved yet.

While previous research on the effect of age on cerebellar gray matter structure used anatomical parcellations, researchers have recently begun exploring novel cerebellar parcellation schemes based on functionally defined regions derived from task-related brain activity (King et al., 2019). The study demonstrated that cerebellar lobular boundaries are poor predictors of functional organization and that functional parcellation have more explanatory power, indicating that anatomical lobules do not reliably correspond to functional subdivisions of the cerebellum. The concept of a functional cerebellar atlas was further advanced by Nettekoven et al. (2025), who integrated data from multiple diverse task-based datasets to produce a more robust and generalizable parcellation. In addition, their atlas adopts a hierarchical organization, enabling the examination of functional regionality at varying levels of detail. At a coarse level, the atlas is divided into four domain-specific regions: motor, action, demand and sociolinguistic. At the medium and fine levels, the cerebellum is divided in 32 and 64 regions, respectively, with each region defined by its functional profile derived from correlations with specific tasks in the dataset. Currently, it remains unclear how age-related structural degeneration manifests within functionally defined regions of the cerebellum.

Next to the typical age-related changes in gray matter volumes, the topological organization of the cerebral cortex, which refers to the way brain regions are arranged and interconnected, has been investigated by looking at inter-individual differences in regional gray matter volumes. While gray matter volume of nearby cortical regions is usually correlated, this is also true for more distant regions such as homologous regions across hemispheres (Mechelli et al., 2005) or regions belonging to the same functional network. For instance, gray matter volume of the hippocampus is correlated to the gray matter volume of other memory areas (Bohbot et al., 2007). Investigating how the topological organization of these networks changes with age can provide us with unique insights about the impact of aging on brain function. For instance, aging has been shown to modify the topological organization of the semantic network and the executive control network (Montembeault et al., 2012). However, it is currently unknown how the topological organization of the cerebellar network changes with age. If some cerebellar subregions are more affected by age than others, one would expect the topological organization of the cerebellum to change with age.

As region-selective vulnerability is characteristic of clinical (cerebellar) disorders (Sarah G Donofrio et al., 2025; Guo et al., 2016; Manto, 2022), this study examines whether age-related cerebellar degeneration also shows regional specificity. To assess the pattern of gray matter loss, we compared regional gray matter volumes between young and older adults for both anatomical and functional cerebellar regions and related topological organization. The inclusion of the functional parcellation rely on the hypothesis that this parcellation may better reflect the function of the cerebellum rather than the anatomical parcellation (Nettekoven et al., 2024) and would make it easier to find a link between structure and function. We expect age-related differentiation across cerebellar regions, with larger age-related differences observed in more anterior anatomical regions (Bernard and Seidler, 2013; Han et al., 2020; Hulst et al., 2015). Likewise, we anticipate differentiation among functionally defined cerebellar regions (Nettekoven et al., 2024), such that age-related differences may be more pronounced in the demand regions, associated with cognitive functions, than in those primarily involved in motor and action processes. These hypotheses were tested across two datasets, one that we collected ourselves (De Witte et al., 2026a) and one that is publicly available (Shafto et al., 2014; Taylor et al., 2017) in order to ensure the generalizability and replicability of our findings.

Further, we examined the relationship between cerebellar regional gray matter volume and sensorimotor performance, using the behavioral data from our previous study (De Witte et al., 2026a). This dataset includes several cerebellar-specific tasks: rhythmic finger tapping (Ivry and Keele, 1989), grip force adjustment (Rost et al., 2005), eye movement coordination (Orban de Xivry et al., 2008, 2006), force matching (Parthasharathy et al., 2020; Wolpe et al., 2016), reach adaptation (Donchin et al., 2012; Smith and Shadmehr, 2005; Vandevoorde and Orban de Xivry, 2021, 2019), speech adaptation (Parrell et al., 2017) and mental rotation (McDougle et al., 2022). Despite minimal age-group differences across most tasks, performance varied widely between individuals. We therefore tested whether this interindividual variability could be explained by regional gray matter volume in anatomically or functionally defined cerebellar regions. We expect that, within the anatomical parcellation, anterior cerebellar regions will account for some variability in sensorimotor performance. However, we hypothesize that motor and action-related regions identified through functional parcellation will provide a stronger explanation of sensorimotor function.

Together, this study aims to clarify how aging affects regional cerebellar gray matter volumes, and how the region-specific cerebellar gray matter organization relates to sensorimotor performance in aging. We further evaluate whether such relationships are better captured by the anatomical or functional organization of the cerebellum.

## 2. Methods

### 2.1 Participants

We recruited 50 young adults between 20 and 25 years and 82 older adults between 55 and 70 years for this study. We also recruited 35 older-old adults above 80 years in the context of a large project on the cerebellum and aging (De Witte et al., 2026b) but this population is not included in the present report as this population had few MR scans (n = 20), which was insufficient for examining within-group variance in gray matter volume and sensorimotor behavior.

Sample size was based on the power needed to detect age-related cerebellar structural changes for a general project on the effect of aging on cerebellar function, using effect sizes from Walhovd et al. (2011); Cohen’s d = 0.98 for YA vs. OA, 0.66 for OA vs. OOA. This required 50 participants per group for the young and older adults to achieve 80% power. Since we also planned to follow the older adult group longitudinally, and wanted to detect correlations around r = 0.3 with 80% power, we needed 65 participants at the second time point. We increased the initial older adult group to 80 to account for an anticipated 20% dropout rate.

During recruitment, participants were verbally screened for right-handedness, non-smoker status and good mental and physical health, including being free of neurological diseases. After providing written informed consent, all participants were screened with the Edinburgh handedness inventory (Oldfield 1971) to confirm their right-handedness. Cognitive functions of older and older-old participants were assessed using the Montreal Cognitive Assessment test (MoCA), for which a minimum score of 23 out of 30 had to be achieved (Carson et al., 2018). Two participants were excluded based on MoCA criteria. The final dataset included a total amount of 50 young (mean age: 23.3 ±2.5, 28 females) and 80 older adults (mean age: 63.1 ±4.6, 45 females). The study was approved by the Ethics Committee Research UZ KU Leuven (project ID: S66650).

### 2.2 Neuroimaging

#### 2.2.1 MRI Acquisition

Images were acquired using a Philips 3T Achieva magnetic resonance scanner (Philips Healthcare, The Netherlands) with a 32-channel receiver head coil, located at the University Hospital of Leuven, Belgium. A high-resolution three-dimensional T1-weighted structural image was acquired (three-dimensional transient field echo (3DTFE); repetition time (TR) = 9.7 ms; echo time (TE) = 4.6 ms; inversion time (TI) = 900 ms; flip angle = 8°; voxel size = 0.89 × 0.89 × 1.0 mm; field of view (FOV)= 256 × 242 × 182 mm; 182 sagittal slices).

T1-weighted imaging data from three participants were missing: from one young and one older participant due to data loss during processing, and from another older participant who was not scanned because MR contraindications were discovered after inclusion.

#### 2.2.2 Cam-CAN imaging data

To compare our cerebellar volume measurements with data from a diffgrerent sample, we made use of structural T1-weighted images from the Cam-CAN repository (Shafto et al., 2014; Taylor et al., 2017). The images were acquired at the Medical Research Council (UK) Cognition and Brain Sciences Unit (MRC-CBSU) in Cambridge, UK, using a Siemens Magnetom TrioTim syngo MR B17 scanner (TR = 2250 ms, TE = 2.99 ms, TI = 900 ms, flip angle = 9°, voxel size = 1.0 × 1.0 × 1.0 mm, FOV = 256 × 240 × 192 mm, multi-slice mode). The dataset included images from 653 participants aged between 18 and 88 years, with approximately 100 individuals per decade.

For comparison with our own study data, we created two subsets of Cam-CAN participants whose ages matched the age groups in our sample: 110 young participants between 20 and 35 years old (mean age 28.8 ± 3.9) and 156 older participants between 55 and 70 years old (mean age: 62.7 ± 4.4).

#### 2.2.3 Cerebellar region analysis

An automated voxel-based morphometry (VBM) analysis was conducted using the Computational Anatomy Toolbox (CAT12.8.2) within SPM12 (Wellcome Trust Centre for Neuroimaging). Analyses of our own images and the Cam-CAN dataset were conducted separately but using similar procedures. For region-of-interest (ROI) volume analysis of the cerebellar anatomically defined regions we used the spatially unbiased atlas template of the cerebellum and brainstem (SUIT) (Diedrichsen, 2006). Prior to the analysis, the atlas was spatially aligned to the shooting template used by CAT12 for segmentation and normalization. Total gray matter volumes for each ROI, as well as total intracranial volume, were directly extracted from the automated VBM analysis. As hemispheric asymmetry was beyond the scope of this study, we summed up left and right hemisphere volumes for each ROI. This resulted in 18 regions: lobules I-IV, lobule V, lobule VI, vermis VI, crus I, vermis crus I, crus II, vermis crus II, lobule VIIb, vermis VIIb, lobule VIIIa, vermis VIIIa, lobule VIIIb, vermis VIIIb, lobule IX, vermis IX, lobule X, and vermis X. A visualization of these regions can be found in Samuelsson et al. (2026). To derive total gray matter volumes for the major lobular divisions of the cerebellum, we summed lobules I–IV and V for the anterior lobe; lobule VI through vermis IX for the posterior lobe; and lobule X and vermis X for the flocculonodular lobe. In parallel, a medial–lateral division separating the vermis from the hemispheric regions was generated. The volumes of these divisions were computed by summing all vermal ROIs and all hemispheric ROIs, respectively.

Based on the same CAT12 segmentation results, a separate ROI analysis was performed, based on the asymmetric functional atlas developed by Nettekoven et al. (2023), which comprises 32 cerebellar regions in MNI152 standard space. To match the non-lateralized approach of the anatomically defined region analysis, left and right hemisphere volumes for each ROI were summed, resulting in 16 bilateral functional cerebellar regions: 4 motor, 3 action, 4 demand, and 5 sociolinguistic regions. To obtain total gray matter volumes for each of the four functional domains, the volumes of all ROIs within each domain were summed. A visualization of this parcellation is available in Nettekoven et al. (2024).

### 2.3 Behavioral assessment

All participants completed approximately five hours of behavioral assessment. For the present study, we selected eight tasks specifically designed to assess cerebellar function: rhythmic finger tapping, grip force adjustment, eye-movement coordination, force matching, reach adaptation, speech adaptation, and mental rotation. Manual tasks were all performed with the dominant right hand. For each task, outcome measures specifically linked to cerebellar processing were isolated from other task-related variables to enable a focused assessment of cerebellar function. We briefly describe each task and the corresponding cerebellar outcome measures below; a comprehensive description is available in de Witte et al. (2026).

#### 2.3.1 Rhythmic finger tapping

Motor timing ability was assessed using a rhythmic finger tapping task (Duchek et al., 1994). Participants were seated at a table with their right index finger on the spacebar of a computer keyboard for tapping in synchrony with an auditory tone sequence (13 tones with 600 ms interval). After the tone sequence stopped, they continued tapping at the same pace for an additional 31 taps. The procedure was repeated across three trials.

For each trial, we used the final 30 inter-tap intervals to calculate clock variance, which reflects variability in the internal timekeeping mechanism and is sensitive to cerebellar function (Wing & Kristofferson, 1973). Trials containing accidental double taps (intervals <150 ms) or abnormally long intervals (>1200 ms, i.e., twice the reference duration) were excluded from the analysis (14 trials in total; 3%). Consequently, four participants (2 young, 2 older) with fewer than two valid trials remaining were excluded.

Clock variances (S_C_) were calculated using the following equations:

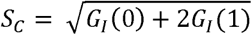

With *G_I_*(1) as the covariance between adjacent inter-tap intervals::

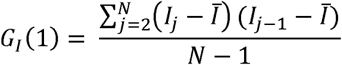

And *G_I_*(0) as total variance of the observable inter-tap intervals:

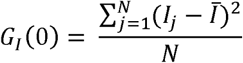

Where *I_J_* refers to the j^th^ response interval, and:

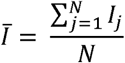

With N = 30 taps. The model predicts that a variation in motor mechanism produces a negative correlation between adjacent intervals. We tested this assumption by calculating whether the lag one serial correlation (PI(1)) was negative, using:

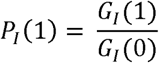

A non-negative *P_I_*(1) value violates the model’s assumptions, suggesting that the expected negative dependency between adjacent intervals (long intervals followed by short ones, and vice versa) is not present. This indicates that all the tapping variability comes from the timekeeper mechanism, and not from motor execution mechanisms, and therefore these trials were retained for analysis (97 trials in total, 26.43%). Additionally, the model assumes that a *P_I_*(1) value below −0.5 points indicates a lack of variability in the timekeeping mechanism. In these trials, the clock variance estimates were set to zero (16 trials in total, 4.36%).

#### 2.3.2 Grip force anticipation

This task assessed the anticipatory modulation of grip force during cyclical object lifting. Participants grasped and moved a lightweight manipulandum vertically between two markers spaced 30 cm apart at a frequency of 2 Hz for a total duration of 10 seconds, corresponding to 10 movement cycles per trial. 5 trial repetitions were performed.

Grip force and load force signals were measured as the forces in the horizontal direction (X) and vertical direction (Z), respectively. Grip force and load force data from the up-down condition were preprocessed to reduce high-frequency noise by applying a fourth-order Butterworth low-pass filter to both, using Matlab’s butter function. A cut-off frequency of 2 Hz was chosen, to exclude noise caused from sources other than the frequency of the object movement. The filter coefficients were determined based on the cutoff frequency and the sampling rate of the data (1000 Hz). The filter was applied in a zero-phase manner using MATLAB’s *filtfilt* function to avoid phase distortion. This two-way filtering process (forward and reverse) ensures the signal remains in phase, allowing accurate temporal alignment of force signals across trials.

To evaluate predictive control, we calculated the average temporal lag between vertical load force peaks and the corresponding grip force peaks across movement cycles. Positive time differences indicate that grip force changes preceded the corresponding load force changes, reflecting anticipatory control. Based on the filtered data, all load force maxima peaks within a 3 s to 9 s time window were identified. Grip force maxima occurring within a 400 ms window centered around each load force peak (−200 ms to +200 ms) were designated as the corresponding grip force peaks. Grip force maxima occurring exactly at ±200 ms from a load force peak were considered unreliable, as they were more likely to represent the highest value at the window boundary rather than a true peak, resulting in the removal of 305 peaks (6.96%). To ensure data quality, participants were required to have at least 10 reliable grip force peaks remaining for inclusion in the analysis, which resulted in the exclusion of one participant. Average time differences between load force and corresponding grip force peaks were calculated to evaluate grip force anticipation.

#### 2.3.3 Eye movement anticipation and coordination

The eye movement coordination task assessed anticipation of eye velocity and the coordination between smooth pursuit and saccadic eye movements during visual tracking of a horizontally moving target (Orban De Xivry et al., 2006). Participants tracked a red dot moving (12 cm/s) at a constant speed on a screen while gaze was recorded using an Eyelink 1000 eye tracker (SR Research Ltd, Kanata, ON) integrated with the Kinarm exoskeleton (Kinarm, BKIN Technologies Ltd., Kingston, ON, Canada). The task included control trials, where the target remained visible throughout its entire motion period (2.4 s), and blanking trials, during which the target was occluded for 1 s midway through its trajectory, requiring participants to maintain tracking of the unseen target.

The experiment started with 20 control trials to familiarize participants with the target trajectory, followed by five test blocks. Each test block comprised six control trials and twenty blanking trials, beginning with two control trials, followed by the remaining trials presented in a pseudorandomized order.

Eye tracking data was filtered with a low-pass, zero-phase autoregressive filter with a cut-off frequency of 20 Hz. Vectorial eye velocity was calculated based on only the horizontal displacement of the eye. To obtain smooth eye velocity, all saccades were removed along with 25 ms of data preceding and following each detected saccade and replaced by linear interpolation. Saccades were identified using an acceleration threshold of 3 cm/s^2^ and a minimum duration of 30 ms.

Eye movement anticipation was quantified as the average smooth eye velocity measured between 50 and 100 ms after target motion onset, based on the control trials. Eye movement coordination was quantified as the regression slope between smooth and saccadic eye movement displacements, expressed as a ratio of total eye displacement. A slope of −1 indicates perfect coordination. Trials containing blinks during the critical periods of interest were excluded from the analysis, resulting in the removal of an average of 4 trials (8.00%) for young adults and 2.39 trials (4.87%) for older adults.

#### 2.3.4 Force matching

The force matching task assessed sensory attenuation during self-applied touch, following the protocol of Wolpe et al. (2016). Participants sat at a custom-built setup consisting of a 10 cm lever attached to a torque motor, with a force sensor measuring forces applied to the left index finger placed underneath the lever. Each trial began with a force perception phase where the torque motor applied a target force (1, 1.5, 2, or 2.5 N) to the participant’s finger for 2.5 seconds. A beep signaled the start of the force reproduction phase, during which participants applied force with their right hand to match the perceived force on their left finger. They verbally confirmed when the forces matched, which was recorded in the data, and held the force for an additional 2.5 seconds until a second beep.

Two reproduction conditions were used: the slider condition (control condition), where participants controlled the torque motor via a slider producing externally generated force, and the button condition, where pressing a button with the right index finger directly transmitted force to the left finger via the torque motor (self-applied force). The experiment comprised 32 trials (eight cycles) per condition, with a short break (±30 s) provided every two cycles. Each task cycle consisted of four trials, with one trial per target force presented in random order.

Both the target and reproduced forces were recorded by a sensor positioned beneath the lever in contact with the participant’s index finger, sampled at 1000 Hz. The reproduced force was calculated as the average self-applied force measured within a 500 ms window centered on the match mark, spanning from 250 ms before to 250 ms after.

Sensory attenuation was quantified as the amount of excess force reproduced during the self-applied force condition relative to the reproduced force in the lever condition. This was calculated by subtracting the average matching error in the slider condition from the matching errors in the button condition.

#### 2.3.5 Reach adaptation

The task-irrelevant clamped feedback paradigm developed by Morehead et al. (2017), was used to assess cerebellar-dependent motor adaptation. Participants performed reaching movements on the Kinarm end-point robot (Kinarm, BKIN Technologies Ltd., Kingston, ON, Canada), from a central home position toward one of four diagonal targets. During the baseline phase (40 trials), visual feedback of hand position was veridical. In the adaptation phase (240 trials), cursor feedback was clamped 30° counterclockwise from the target, independent of actual hand direction. Participants were informed of this manipulation and instructed to ignore the cursor motion and to continue reaching directly toward the target. Hand error was defined as the angular deviation between the hand and target directions, measured 4 cm into the reach. Trials with a hand error exceeding ±60° were excluded, as such large deviations typically reflected inaccurate reaching caused by predicting the target at an incorrect location, resulting in the removal of 444 trials in total (1.25%) from the dataset. Final implicit adaptation was calculated as the average change in hand error over the last 40 adaptation trials, corrected for individual baseline error.

#### 2.3.6 Speech adaptation

We assessed speech adaptation using an auditory feedback perturbation paradigm adapted from Parrell et al. (2017). Participants read aloud the word “bed” while hearing real-time auditory feedback of their own voice through headphones. During the 30-trial baseline phase, this feedback was unaltered. In the 90-trial adaptation phase, the first vowel formant (F1) was shifted by –125 Mels, making “bed” sound more like “bid.” This manipulation was introduced abruptly and without the participant’s awareness. Speech was recorded and processed in real time. Vowels were extracted from sound peaks exceeding an amplitude threshold of 0.025, or 0.04 for participants whose vowel peaks were not captured by the lower threshold upon visual inspection. F1 values were extracted from the vowels using analysis software from Parrell et al. (2017). Trials were excluded based on the following criteria: short vowel duration (<120 ms); extremely low (<250 Mels) or high (>1000 Mels) F1 frequencies indicating incorrect vowel tracking; or F1 values deviating by more than three standard deviations from most trials. In total, 478 trials (2.92%) were excluded. A baseline F1 reference was calculated as the median F1 across the last 25 baseline trials. Final F1 adaptation was quantified as the change in F1 across the last 30 adaptation trials, normalized by subtracting the baseline reference.

#### 2.3.7 Mental rotation

We assessed cerebellar contributions to internal model-based spatial transformations using a mental rotation task adapted from McDougle et al. (2020). Participants viewed white capital letters (F, G, J, R and their mirror-reflected versions) presented in various rotated orientations (±15°, ±75°, ±135°) and judged whether each letter was normal or mirrored by pressing a corresponding key of a computer keyboard. Stimuli remained on the screen until a response was made, and accuracy feedback was provided after each trial. Stimuli were presented in a random order, with an equal number of normal and mirrored presentations of each stimulus at each rotation sign and magnitude. After each trial, feedback indicating whether the response was correct was displayed on the screen. The task consisted of 144 trials, with an inter-trial interval of 2 s.

Only trials with correct responses were included in the analysis, comprising 93.21% of trials for young adults and 93.53% for older adults. Based on these trials, a regression line was fitted through the mean reaction times at target rotations of 0°, 15°, 75°, and 135. The slope of this line, reflecting the increase in response time per degree of rotation, was taken as a measure of mental rotation pace.

### 2.4 Data analysis

#### 2.4.1 Regional gray matter differences between age groups

To assess the effects of age group on cerebellar gray matter volume while controlling for the total intracranial volume, a multivariate analysis of covariance (MANCOVA) was performed. Adding the total intracranial volume as a covariate rather than normalizing all the gray matter values by this value was preferred as it is often more powerful (Sanfilipo et al., 2004). For analysis of the anatomically defined cerebellar regions, the dependent variables were the gray matter volumes from the 18 anatomically defined regions: lobules I-IV, lobule V, lobule VI, vermis VI, crus I, vermis crus I, crus II, vermis crus II, lobule VIIb, vermis VIIb, lobule VIIIa, vermis VIIIa, lobule VIIIb, vermis VIIIb, lobule IX, vermis IX, lobule X, and vermis X. In addition, we conducted separate analyses to examine age-group effects on the total gray matter volume in two additional anatomical subdivisions: the anterior, posterior, and flocculonodular cerebellar lobes (lobular division), as well as on the vermal and hemispheric regions (medial-lateral division). For the analysis of the functionally defined cerebellar regions, a separate analysis was performed to determine the gray matter volume of 16 distinct cerebellar regions based on a functionally-informed segmentation scheme (Nettekoven et al., 2024): 4 motor regions, 3 action regions, 4 demand regions and 5 sociolinguistic regions. In addition, we examined age-group effects across the functional domains to which the regions belonged (motor, action, demanding, and sociolinguistic).

For both the anatomically and functionally defined region analyses, MANCOVA was conducted using MATLAB’s *mancovan* toolbox, using the Lawley-Hotelling trace statistic to evaluate multivariate effects, including total intracranial volume as a covariate. When the MANCOVA indicated a significant group effect, post hoc univariate ANCOVAs were performed for each ROI to determine which specific regions contributed to the overall group effect. For each region, the F-statistic and p-value were extracted. To correct for multiple comparisons across the regions, false discovery rate correction was applied using the Benjamini–Hochberg procedure.

For effect sizes of post-hoc comparisons, Cohen’s d was calculated based on F-values of between-group effects, calculated as:

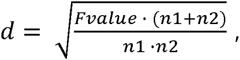

in which n is the group size. Corresponding confidence intervals were computed based on the standard error of d, using 95% confidence.

We tested whether age-group effects varied across anatomically and functionally defined regions, as well as across the anatomical subdivisions (lobular and medial-lateral) and functional domains by fitting separate mixed-effects models. The applied models were specified as

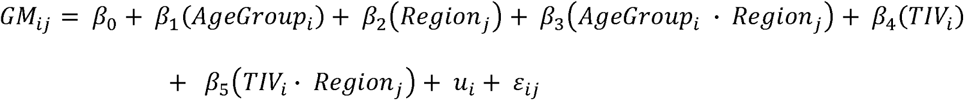

Here, gray matter volume (*GM*) for participant *i* and region *j* served as the dependent variable. *β*_0_ is the intercept representing baseline gray matter volume at reference levels of *AgeGroup* and *Subregion*. *β*_1_ and *β*_2_ capture the fixed effects of age group and cerebellar division, respectively. *β*_3_ is the interaction term testing whether age-group effects differ between divisions. *β*_4_ is the covariate effect of total intracranial volume and *β*_5_ captures the fact that all regions are differently related to TIV. Age group and region were entered as categorical variables: young adults coded as 1 and older adults as 2; regions were numbered by region (e.g., Region1, Region2, etc.). Because multiple regions were measured within each participant, we included a random intercept for participant (*u_i_*), allowing individual baseline differences and accounting for within-participant correlation. The residual error is represented by *ε_ij_*. Significance of fixed effects, including the Age Group x Division interaction, was assessed using Satterthwaite-adjusted degrees of freedom.

Using the matched age groups from the Cam-CAN dataset, we conducted analogous analyses across the anatomically and functionally defined cerebellar regions as well as the lobular, medial-lateral divisions and functional domains. In addition to this cross-sectional comparison between young and older adults, we analyzed the full Cam-CAN cohort to assess the linear relationship between gray matter volume and age (Supplementary methods).

#### 2.4.2 Gray matter correlations between regions

To characterize structural covariance within the cerebellum, we quantified inter-regional correlations in gray-matter volume across cerebellar regions. This approach allows us to identify which regions show stronger versus weaker structural coupling. We computed partial correlation coefficients between regional gray-matter volumes while controlling for total intracranial volume. Prior to analysis, both regional gray-matter volumes and total intracranial volume were robustly z-scored within each age group. Partial correlation matrices were then generated for the full sample and separately for each age group, providing insight into overall covariance patterns as well as potential age-related differences in structural coupling.

For the Cam-CAN dataset, we generated similar partial correlation matrices: one including all participants and two corresponding to the defined age groups. In the analysis including all participants between 18 to 80 years old from the Cam-CAN dataset we could not perform z-scoring for each age group (as age was a continuous variable). To make full use of the broad age range, we therefore included age as an additional covariate when computing the partial correlation coefficients.

To statistically examine which inter-regional correlations differed in strength between the two age groups from our own data and from the Cam-CAN dataset, we estimated between-group differences for each pairwise regional correlation using a non-parametric bootstrap procedure (N=5000). In each iteration, participants from both groups were resampled with replacement, and for each group a new partial correlation matrix was computed. For each resampled dataset, correlation differences between the matrices of young and older participants were calculated, generating a null distribution. From this distribution, we derived two-tailed p-values and 95% confidence intervals per cerebellar region, defined as the proportion of bootstrap samples in which the absolute correlation difference exceeded the observed difference.

For the functionally defined cerebellar regions, the partial correlation matrices revealed stronger correlations among regions within the same functional domain than among regions belonging to different domains. Because age-group differences in these patterns were minimal, we conducted this analysis on the combined dataset of both age groups. For within-domain correlations, we calculated the mean partial correlation coefficient for each domain, based on all pairwise partial correlations among regions belonging to that domain. For between-domain correlations, we computed the average partial correlation coefficient across all region pairs spanning two distinct domains. This resulted in 12 unique domain-level correlation estimates. To compare within-domain correlations with between-domain correlations, we performed another non-parametric bootstrap (N = 5000 iterations) that calculated the differences between those for each of the domains separately based on the resampled data.

#### 2.4.3 Factors accounting for inter-regional correlation coefficients

##### 2.4.3.1 Anatomically defined regions

Based on the gray matter partial correlation coefficients between regions, we examined whether these values could be predicted by a subset of anatomical and spatial factors. For correlations between anatomically defined regions, we included three predicting factors: (1) the spatial proximity between regions, (2) whether the regions belonged to the same lobular division (anterior, posterior, or flocculonodular), and (3) whether they were part of the same medial–lateral division (vermis or hemispheres). Spatial proximity was quantified by computing the mean Euclidean distance between the centers of gravity of each region (Supplementary results Table 1). Because both matrices were symmetric, we extracted and vectorized the upper triangular portions, which were then used as predictors of the observed partial correlations. To determine to what extent the observed correlations between regions could be explained by the predictors, we conducted robust multiple linear regression analyses. The dependent variables were the partial correlation matrices, while the independent variables were the predictor matrices. Four models were fitted:

1. a full model including all predictors:

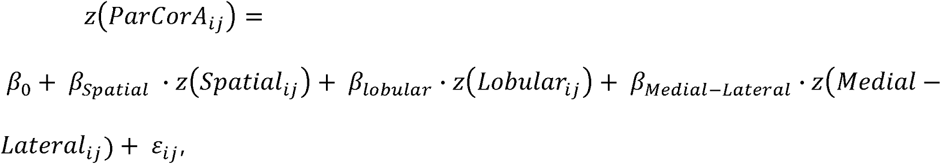
2. a spatial-only model including spatial distance only

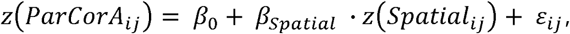
3. a lobular-only model, including anterior-posterior-flocculonodular division only

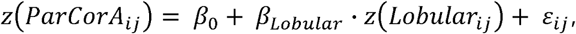
4. a medial-lateral-only model, including vermis-hemisphere division only

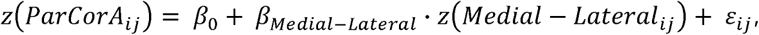

In these equations *z*(*ParCor_ij_*) is the z-scored partial correlation between regions i and j, controlling for total intracranial volume (and age when considering all the participants from the Cam-CAN dataset). *Z*(*Spatial_ij_*) is the z-scored Euclidean distance between regional centers of gravity, *Z*(*Lobular_ij_*) is the z-scored lobular division between anterior, posterior, and flocculonodular regions, *Z*(Medial − *Lateral_ij_*) is the z-scored medial-lateral division between vermis and hemisphere regions. For both lobular and medial-lateral divisions this is 1 for regions of the same division and zero for regions from different divisions. *β*_0_, is the intercept, *β_Spatial_*, *β_Lobukar_*, and *β_Medial−Lateral_* are robust regression coefficients estimated with Matlab’s *robustfit* function, and *ε_ij_* is the residual error.

We computed the coefficient of determination (R^2^) for each model to assess explanatory power. To estimate the stability and statistical significance of the regression coefficients and model fits, we performed a bootstrap analysis with 5000 iterations. In each iteration, a resampled subset of participants was used to recompute the partial correlation matrix and repeat the regression analysis (regression coefficients and R^2^). Confidence intervals (95%) were calculated for each regression coefficient and R^2^ value, by taking the 2.5th and 97.5th percentiles of the bootstrap distribution, corresponding to the lower and upper bounds of the interval. Two-tailed p-values were estimated to assess differences in R^2^ between each pair of models, by computing the proportion of bootstrap iterations where one model outperformed the other.

##### 2.4.3.2 Functionally defined regions

A similar predictive model was constructed to explain variance in the partial correlation coefficients across functional regions. In this model, we tested the extent to which higher inter-regional correlations could be accounted for by spatial proximity or by functional similarity. Functional similarity was quantified by computing correlation coefficients between regional functional profiles derived from the Multi-Domain Task battery reported by Nettekoven et al. (2024). Spatial proximity between functional regions was estimated using the centers of gravity of their respective compartments, as defined in the 128-region version of the Nettekoven atlas. To approximate inter-regional distance as accurately as possible, we calculated the mean Euclidean distance between all compartment centers of gravity in one region and all compartments of the other region. Three models were fitted:

1. a full model including both functional similarity and spatial distance:

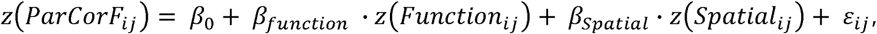
2. a function-only model including functional similarity only

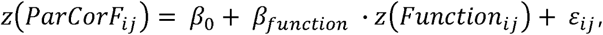
3. a spatial-only model, including spatial distance only

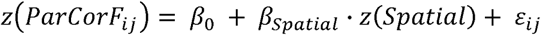

In these equations *z*(*ParCor_ij_*) is the z-scored partial correlation between regions i and j, controlling for total intracranial volume (and age when considering all the participants from the Cam-CAN dataset). *Z*(*Function_ij_*) is the z-scored functional similarity where *Function_ij_* is equal to one if both regions are part of the same domain (motor, action, demand, or sociolinguistic) and zero otherwise. *Z*(*Spatial_ij_*) is the z-scored Euclidean distance between regional centers of mass.*β*_0_ is the intercept, *β_Function_* and *β_Spatial_* are robust regression coefficients estimated with Matlab’s *robustfit* function, and *ε_ij_* is the residual error. For the comparison of these models, we also calculated p-values and corresponding 95% confidence intervals to assess differences in performance.

#### 2.4.4 Correlations between behavioral tasks

If age-related degeneration of the cerebellum directly impacts function, we should expect that cerebellar outcomes from tasks that involve the same effector or from the same functional domain would be correlated with each other, especially in older participants. To test this, we calculated the correlations between the results of the different behavioral tasks to determine whether there was a general pattern of performance across tasks, or whether performance was more task-specific. First, we included all participants to assess general performance patterns. Additionally, correlations were computed separately for each age group to examine whether these patterns differed between younger and older participants.

Before doing so, behavioral task outcome measures were normalized using robust z-scores computed within each age group. For each variable, the robust z-score was defined as:

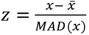, where x̄ is the median value within the corresponding age group, and MAD is the unscaled median absolute deviation (i.e. the median of the absolute deviations from the group median). This approach avoids the influence of outliers while maintaining comparability within groups. Missing values were excluded from the computation of medians and MADs. Pairwise Spearman correlations were computed between robust z-scored task outcomes. This combined approach, using rank-based correlation on robustly normalized data, provides resistance to outliers and non-normal distributions, ensuring more stable estimates of association in heterogeneous data.

#### 2.4.5 Predicting task performance from regional gray matter volume

To test whether cerebellar regional gray matter volume could predict task performance, we constructed two regression models per task: one for the anatomically defined regions and one for the functionally defined regions. Each of the models were performed on the full sample, and additionally within each age group, with age-specific results reported in the Supplementary Results (Supplementary Fig. 3).

For each task, a noise ceiling was calculated to estimate the maximum explainable variance. Noise ceilings were derived using a split-half reliability approach, in which the outcome measure of interest was averaged separately for odd and even trials within each participant. The correlation between these two estimates, corrected using the Spearman-Brown formula, was taken as the maximum explainable variance for that task (i.e. the noise ceiling; Van Bree et al., 2025). Noise ceilings for the full-sample models were computed using performance data from all participants, whereas noise ceilings for the age-group– specific models were based solely on data from the corresponding age group.

Behavioral measures and gray matter data used for the models were standardized within age groups using a robust z-scoring procedure. Gray matter volumes were normalized relative to total intracranial volume before z-scoring, to control for global brain size differences. Since the obtained gray matter partial correlation matrices indicated high inter-regional gray matter correlations in both anatomically and functionally defined regions (Fig. 2A), we used ridge regression (MATLAB’s *ridge function*) to estimate model coefficients. We selected a relatively low regularization parameter (λ = 1) to minimize coefficient shrinkage while still accounting for multicollinearity among predictors.

A leave-one-out cross-validation approach was applied to assess the predictive validity of each model. For each left-out participant, ridge regression was trained on the remaining participants and used to predict the excluded participant’s performance. Cross-validated values were computed from the total sum of squares and residual sum of squares of leave-one-out cross-validation predictions.

To estimate the stability and significance of model coefficients, 5,000 bootstrap resamples were generated per task. On each bootstrap sample, ridge regression coefficients were re-estimated. From the bootstrap distributions, we computed the mean coefficient estimates and two-sided bootstrap p-values.

## 3. Results

We assessed gray-matter volumes from different cerebellar regions in 50 young and 80 older adults collected in our laboratory and from the Cam-CAN dataset using both a classical anatomical parcellation based on cerebellar lobules and a more recent parcellation approach that defines cerebellar regions according to functional specificity (Nettekoven et al., 2024). With these datasets, we investigated age-related changes in the structure of the cerebellum and in its topological organization via the structural covariance matrix. Finally, we investigated how the structure can predict the function based on the behavioral outcomes from eight tasks that probed different aspects of cerebellar sensorimotor function, including motor timing, anticipatory grip control, eye-movement coordination, sensory attenuation, motor adaptation, speech adaptation, and mental rotation.

### 3.1 Age-related effect on cerebellar gray matter is heterogeneous across anatomically and functionally defined cerebellar regions

#### 3.1.1 Anatomically defined cerebellar regions

We assessed gray matter differences between young and older adults across the 18 anatomically defined cerebellar regions and found a strong multivariate effect of age group on regional gray matter, controlled for total intracranial volume (effect of group: Lawley-Hotelling trace *T* = 92.48, *p* < 0.001). Total intracranial volume, included as a covariate, showed a positive correlation with gray matter volume across all regions (main effect of total intracranial volume: Lawley–Hotelling trace T = 244.4, p < 0.001). In addition, we found evidence that the effect of age differed across anatomically defined regions (Fig. 1A, orange data: region × age-group interaction: F(17, 2159) = 18.82, p <0.001). While Lobule VI (d=1.12), lobule V (d=0.94) and Crus I (d=0.92) exhibited the largest age effects, Vermis X (d=0.016), Vermis Crus II (d=0.13), Lobule IX (d=0.28), Lobule VIIIa (d=0.31) or Lobule VIIIb (d=0.35) showed very small (and non-significant) variations across the two age groups.

**Figure 1.**
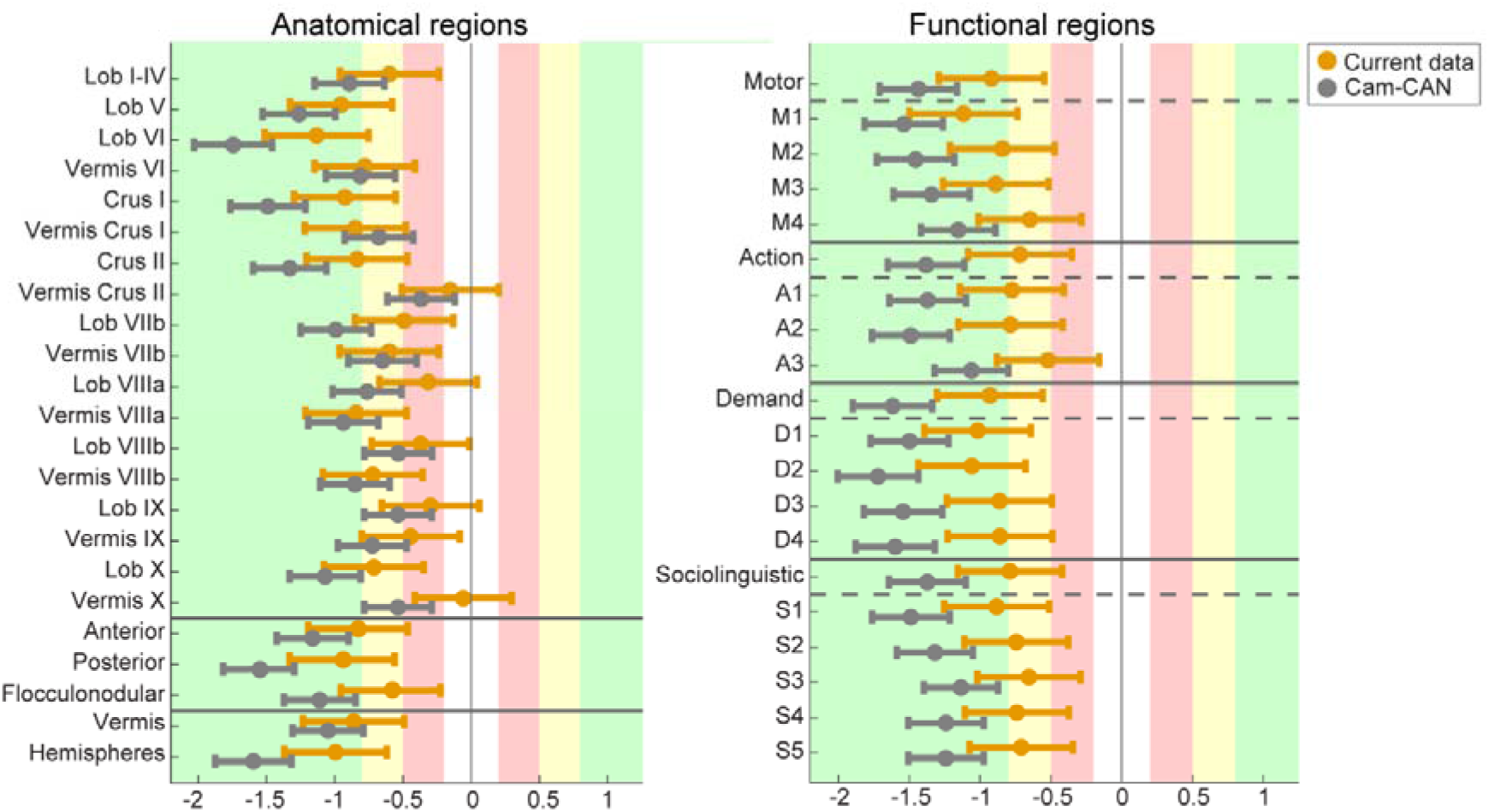
Overview of effect sizes for gray matter volumes in older adults versus young adults. Each dot represents the Cohen’s d effect size for the comparison between young and older adults for each cerebellar region. Data from our dataset are shown in orange, and data from the Cam-CAN cohort are shown in gray. Horizontal error bars indicate 95% confidence intervals. Positive values reflect larger gray matter volumes in older adults relative to young adults, whereas negative values indicate smaller volumes. Cohen’s d values can be interpreted as: negligible (<0.2), small (0.2–0.5), medium (0.5–0.8), and large (>0.8). **(A)** Effect sizes shown for the anatomical division, first grouped by specific cerebellar regions, then by major anatomical divisions (anterior, posterior, and flocculonodular lobes), and finally by vermis versus hemispheric regions. The anatomical divisions are separated by solid gray horizontal lines. **(B)** Effect sizes are shown for the functional divisions of the cerebellum. Within each functional domain, the domain-level effect size is displayed first, followed by the effect sizes of its constituent subregions, which appear below the dashed gray horizontal line. Different functional domains are separated by solid gray horizontal lines.

To evaluate whether the age differences on regional cerebellar gray matter volume in our sample align with those in a larger reference population, we additionally analyzed gray matter volume in anatomically defined regions using images from the Cam-CAN dataset (Shafto et al., 2014; Taylor et al., 2017). Because Cam-CAN includes a continuous age range, we created two age groups that matched those in our study: a young adult group (20-35 years old, n = 110, mean age = 28.84 ± 3.94) and an older adult group (55-70 years old, n = 156, mean age = 62.74 ± 4.44). Consistent with findings in our dataset, a significant multivariate effect of age on regional gray matter volume, controlled for intracranial volume, was found (Lawley-Hotelling trace T = 311.37, p < 0.001). A positive effect of intracranial volume on regional gray matter volume was found in the Cam-CAN data as well (Fig. 1A gray data: Lawley-Hotelling trace T = 324.46, p < 0.001). In addition, age effects were heterogeneous across anatomically-defined cerebellar regions (interaction between age groups and regions: F(17,4522)= 99.54, p<0.001). The same three regions as in our own dataset showed the largest age effects: Lobule VI (d=1.74), lobule V (d=0.1.26) and Crus I (d=1.49). Compared to our dataset, the same set of regions showed the smallest age effects: Vermis X (d=0.54), Vermis Crus II (d=0.37), Lobule IX (d=0.54), and Lobule VIIIb (d=0.54). To assess continuous age differences on gray matter volume in each anatomically defined region, we additionally estimated regression slopes using the full Cam-CAN sample (ages 18–88 years; n = 653). In this case, we also found that the relationship between age and grey matter volumes corrected for total intracranial volume differed across anatomical cerebellar regions (F(17,11101)=339, p<0.001). These results are presented in the Supplementary Results (Supplementary Figure 1).

Motivated by evidence from previous studies that age-related degeneration is most pronounced in the anterior cerebellum and limited to distinct regions within the posterior lobes (Bernard and Seidler, 2013; Hulst et al., 2015), we used the lobular division of the cerebellum to assess age differences on gray matter volume in the anterior, posterior, and flocculonodular divisions. We found a main effect of age on gray matter volume in these divisions (Lawley-Hotelling trace T = 30.9, p < 0.001). The age effect on gray matter volume varied across these subdivisions (Fig. 1A): the anterior part (d = 0.83), posterior (d = 0.95), and flocculonodular divisions (d = 0.6). The magnitude of this effect differed statistically across these divisions (interaction between lobular division and age: F(2,254) = 26.25, p<0.001). Separate ANOVA’s for each pair of subdivisions revealed a significant interaction in all cases (interaction between regions and age group: anterior vs. posterior: F(1,127)= 24.75, p<0.001; anterior vs. flocculonodular: F(1,127)=19.2, p<0.001; posterior vs. flocculonodular: F(1,127)=27.7, p<0.001).

A comparable age effect was found in the Cam-CAN data (Lawley-Hotelling trace = 173.11, p < 0.001), showing similarly pronounced effects across the anterior (d = 1.16), posterior (1.57), and flocculonodular divisions (d = 1.11). There is evidence for a differences across divisions (interaction between lobular division and age: F(2,532) = 153.7, p < 0.001).

Lastly, we divided the cerebellum into vermis and hemispherical regions. In our dataset, age group significantly affected gray matter volume in these regions (Lawley– Hotelling trace = 29,24) with the magnitude of age-related differences significantly varying between regions (interaction between hemispheric division and age: F(1, 127) = 29.48, p < 0.001). The strongest age-related effect was found in the hemispheres (d = 0.98) compared to the vermis (d = 0.83), confirming that the effect differed across these regions. This age effect was also observed in the Cam-CAN data (Lawley–Hotelling trace = 165.13, p < 0.001), also showing a difference in age-related effects between hemispheres (d = 1.6) and vermis volumes (d = 1.05) (interaction between hemispheric division and age: F(1,266) = 166.85, p < 0.001).

#### 3.1.2 Functionally defined regions

In addition to the anatomically defined cerebellar regions, we examined gray matter differences between young and older adults across the four domains and across the 16 functionally defined cerebellar regions based on the atlas developed by Nettekoven et al. (2024). When assessing age differences in each of the functional domains, all of the four domain regions showed smaller gray matter volumes in older adults compared to young adults. In our dataset, the magnitude of the effect differed across the four domains (interaction between age and functional domains: F(3,381) = 9.2, p <0.001). The age effect was larger for the motor and demand domains (d=0.9 and d=0.92, resp.) than for the action and sociolinguistic domains (d=0.7 and d=0.78, resp.). Comparing age-related differences for each pair of domains separately, we found age-related differences in gray matter volume were smaller in the action domain than in the other domains (interaction between age and a pair of domains: all p-values <0.001; motor vs. action: d=0.95; demands vs. action: d=0.91; sociolinguistic vs. action: d=0.66). For the three other comparisons across domains, there was no evidence for a differential effect of age across them (motor vs. demands: d=0.21, p=0.25; motor vs sociolinguistic: d= 0.2, p=0.28; demand vs. sociolinguistic: d= 0.08, p=0.68).

This result was confirmed in the Cam-CAN dataset, which also provided evidence for differences across age groups between the four domains (interaction between age and domain: F(3,798) = 60.08, p < 0.001). However, the effect sizes were larger and the relative changes across the different domains differed compared to our own dataset. The age-related difference in gray matter was still the largest for the demand domain (d = 1.62), followed by the motor domain (d=1.44) with the action and sociolinguistic domains following closely (d= 1.38 and d=1.37). When comparing the gray matter volumes for each pair of domains, separately, five of the six interactions between age group and regions were significant (all p-values<0.005; action vs. motor: d=1.21; action vs. demand: d=1.5; action vs. sociolinguistic: d=1.1; motor vs. demand: d=0.57; motor vs. sociolinguistic: d=0.35). The age-related difference in volume did not differ between the demand and sociolinguistic domains (d=0.2, p=0.09). Analysis of the whole Cam-CAN dataset confirmed this observation as the slope of the relationship between age and grey matter varied across the four domains (F(3,1959)=230.37, p<0.001). The slope of the relationship between age and grey matter volume corrected for total intracranial volume was the steepest for the Demand domain (−3.9% per decade, CI = [−4.2 −3.6]), followed by the Sociolinguistic domain (−3.5% per decade, CI = [−3.8 −3.2]) and by the action and motor domains (for both: −3.2% per decade, CI = [−3.6 −2.8]).

Similar to what we found for the four domains, we found a multivariate effect of age group on regional gray matter volume of the 16 functionally defined regions, controlling for total intracranial volume (Lawley-Hotelling trace *T* = 101.42, *p* < 0.001). Total intracranial volume positively affected regional gray matter volume (Lawley–Hotelling trace *T* = 292.63, *p* < 0.001). The age effect differed across the 16 functional regions (**Error! Reference source not found.**B, orange data: interaction between functional regions and age: F(15,1905) = 10.5, p < 0.001). The variability of this effect is displayed in Fig.1B. Results from the Cam-CAN analysis confirmed the existence of a significant age effect on gray matter volumes in the functionally defined regions, while controlling for total intracranial volume (Lawley– Hotelling trace T = 290.67, p < 0.001). This was also true when controlling for the effect of intracranial volume on regional gray matter volume (Lawley-Hotelling trace T = 316.96, p < 0.001). The Cam-CAN dataset also confirmed that age effects varied across the 16 functionally defined regions (F(15,3990)=76.93, p<0.001). The regression slopes between gray matter volume and age for each functional region, based on the full Cam-CAN sample are shown in the Supplementary Results (Supplementary Figure 2). This analysis also confirmed that the relationship between age and grey matter volume varied across the 16 regions (F(15,9795)=274.52, p<0.001).

### 3.2 Topological organization of the cerebellum is not affected by aging in both datasets

#### 3.2.1 Anatomically defined regions

We examined the extent to which gray matter volumes of different cerebellar regions correlate with each other to determine whether they follow a general pattern across individuals or reflect region-specific variations. We examined inter-regional correlations of gray matter volume controlled for total intracranial volume across the entire sample. We observed that partial correlation were strongest among anatomically adjacent regions, as indicated by higher coefficients along the diagonal of the correlation matrix (Figure 2A, left). This pattern was also present in both younger and older adults when the sample was stratified by age (Figure 2A, middle and right).

**Figure 2.**
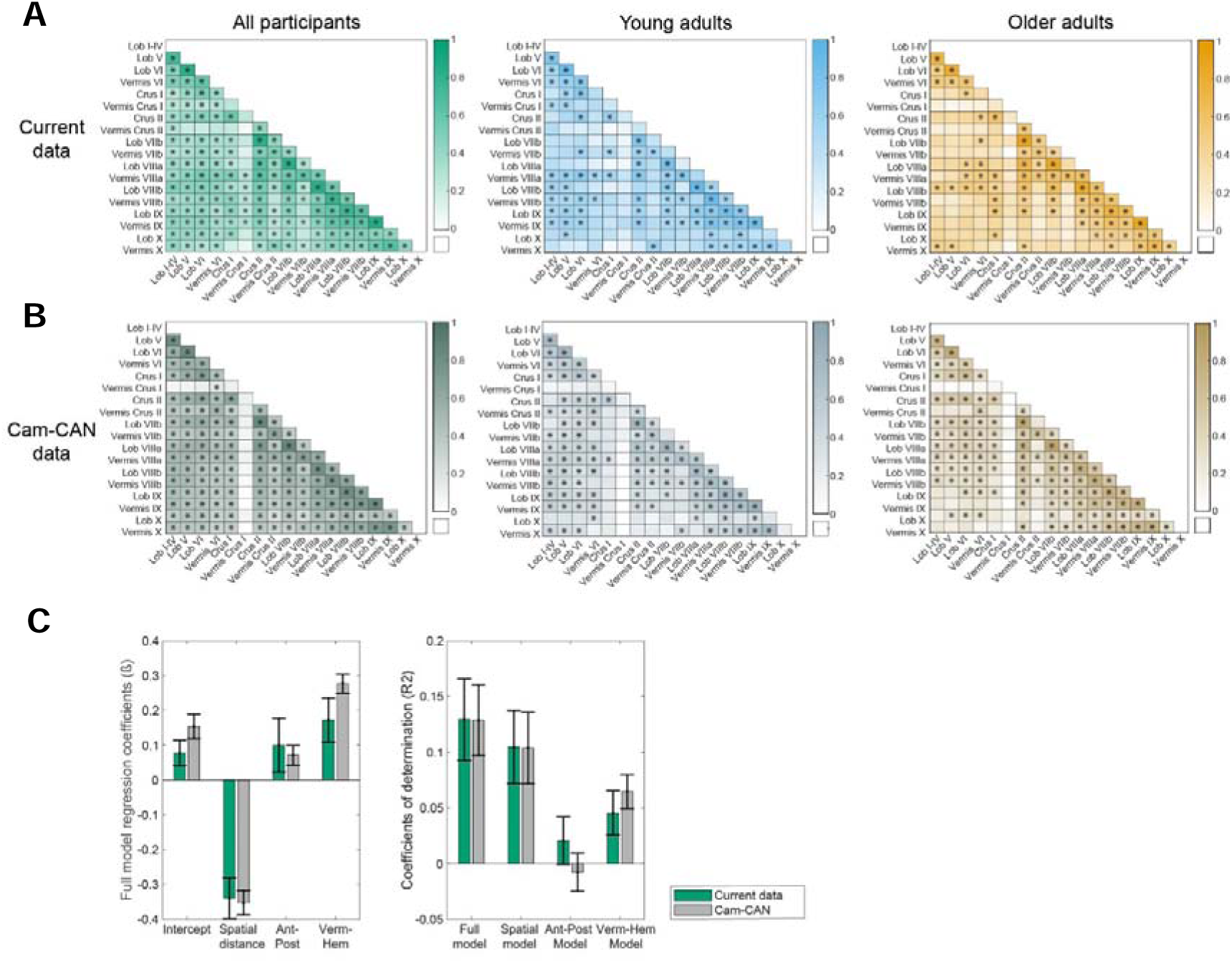
Correlations between anatomical cerebellar regions. **(A)** Mutual gray matter volume partial correlations between anatomical cerebellar regions in our dataset, shown for all participants combined (left, green), young adults only (middle, blue), and older adults only (right, orange). Higher correlations appear in more saturated colors, whereas lower correlations appear lighter. Significant correlations are marked with a star. **(B)** Mutual gray matter volume correlations between anatomical cerebellar regions in the Cam-CAN, shown for all participants combined (left, gray-green), young adults only (middle, gray-blue), and older adults only (right, gray-orange). **(C)** Left: Bar plots showing the mean beta coefficients from the full model predicting gray matter correlations between regions as a function of spatial distance, lobular division, and medial–lateral division, for our own dataset (green) and the Cam-CAN dataset (gray). Error bars represent standard deviations. Right: Bar plots showing the mean R^2^ values for the full model, which includes all predictors, and for the reduced models that include each predictor individually, for our own dataset (green) and the Cam-CAN dataset (gray). Error bars represent standard deviations

When investigating differences in correlation patterns between young and older adults, primarily vermal region correlations seemed to be reduced in older adults, as well as the correlation between crus I and lobule VI. However none of these age-related differences in correlations remained significant after applying the Benjamini-Hochberg correction (p > 0.05).

In the Cam-CAN dataset, we observed similar patterns of stronger correlations between adjacent regions (Figure 2B). In this dataset, Vermis Crus I deviated from the otherwise consistent pattern of inter-regional correlations, showing minimal association with other cerebellar subregions. Notably, Vermis Crus I was by far the smallest region in the dataset (mean volume: 8.5 mm^3^, compared to next smallest region: 145.0 mm^3^), which may have reduced measurement reliability and statistical power, potentially attenuating observable correlations. No correlation differences were found between young and older adults in the Cam-CAN dataset (p > 0.05, Benjamini-Hochberg corrected).

To determine whether the observation that spatially closer regions exhibit higher gray-matter correlations could be statistically substantiated, we evaluated the extent to which inter-regional correlations were predicted by spatial proximity. We further examined whether these correlations were better accounted for by anatomically defined factors, specifically whether regions belonged to the same lobular division (anterior, posterior, or flocculonodular). To do so, we performed a robust multiple linear regression, using the partial correlation coefficients as the dependent variable, the Euclidean distance between regions, and their membership to the same lobular or medio-lateral region as predictors. The full model included all three predictors and explained, in our own data, a small portion of the variance in partial correlation (R^2^_full_ = 0.13). Spatial distance contributed significantly (ß_spatial_ = −0.34, p < 0.001), as did vermis-hemisphere grouping (ß_Medial-Lateral_ = 0.17, p = 0.008). These results confirm that regions in closer spatial proximity show stronger correlations and further indicate that correlations are higher when both regions belong to the vermis or both are located within the cerebellar hemispheres. Conversely, anterior-posterior grouping did not show a significant effect (ß_Lobular_ = 0.10, p = 0.19; Figure 2C).

To assess the relative contributions of each predictor, we compared the full model to simpler models using bootstrap resampling (*N* = 5000). These models included a spatial model (spatial distance as the sole predictor), a lobular model (anterior–posterior grouping as the sole predictor), and a medial-lateral model (vermis–hemisphere grouping as the sole predictor). The full model performed significantly better than each of the reduced models (for all comparisons: p < 0.05; R^2^_spatial_ = 0.10, R^2^_Lobular_ = 0.02, R^2^_Medial-Lateral_ = 0.05; Figure 2C, right). Additionally, the spatial model performed better than the lobular model (p = 0.005), but not better than the medial-lateral model (p = 0.11). No difference in performance was found between the anterior-posterior-only model and the vermis-hemisphere-only model (p = 0.37). These results indicate that gray matter volume correlations between anatomically regions are best explained by spatial proximity and anatomical grouping according to vermis or hemispheric location, although together these factors account for only a small portion of the variance.

These findings were replicated in the Cam-CAN dataset in which the full model explained a comparable proportion of variance (R^2^_full_ = 0.12), with all predictors contributing significantly (ß_spatial_ = −0.35, p < 0.001; ß_Lobular_ = 0.07, p = 0.01; ß_Medial-Lateral_ = 0.28, p < 0.001). Moreover, the full model outperformed each of the simpler models (all comparisons: p < 0.05; R^2^_spatial-only_ = 0.10, R^2^_Lobular_= =0.007, R^2^_Medial-Lateral_ = 0.06). Finally, the spatial model performed significantly better than the lobular model (p < 0.001), but not better than the medial-lateral model (p = 0.10).

#### 3.2.2 Functionally defined regions

We examined the gray matter volume correlations between functionally defined cerebellar regions. When analyzing all participants together, positive associations were observed between all pairs of regions, with particularly strong associations between regions from the same functional domain (all correlations: r > 0.53, *p* < 0.001; Figure 3A, left). When analyzing both groups separately, positive associations between regions were found in both age groups (r_Young_ > 0.46; r_Older_ > 0.53), showing highly similar correlation patterns (Figure 3A, middle and right). When statistically comparing correlation coefficients between young and older adults we did not find any differences that survived the Benjamini-Hochberg correction.

**Figure 3.**
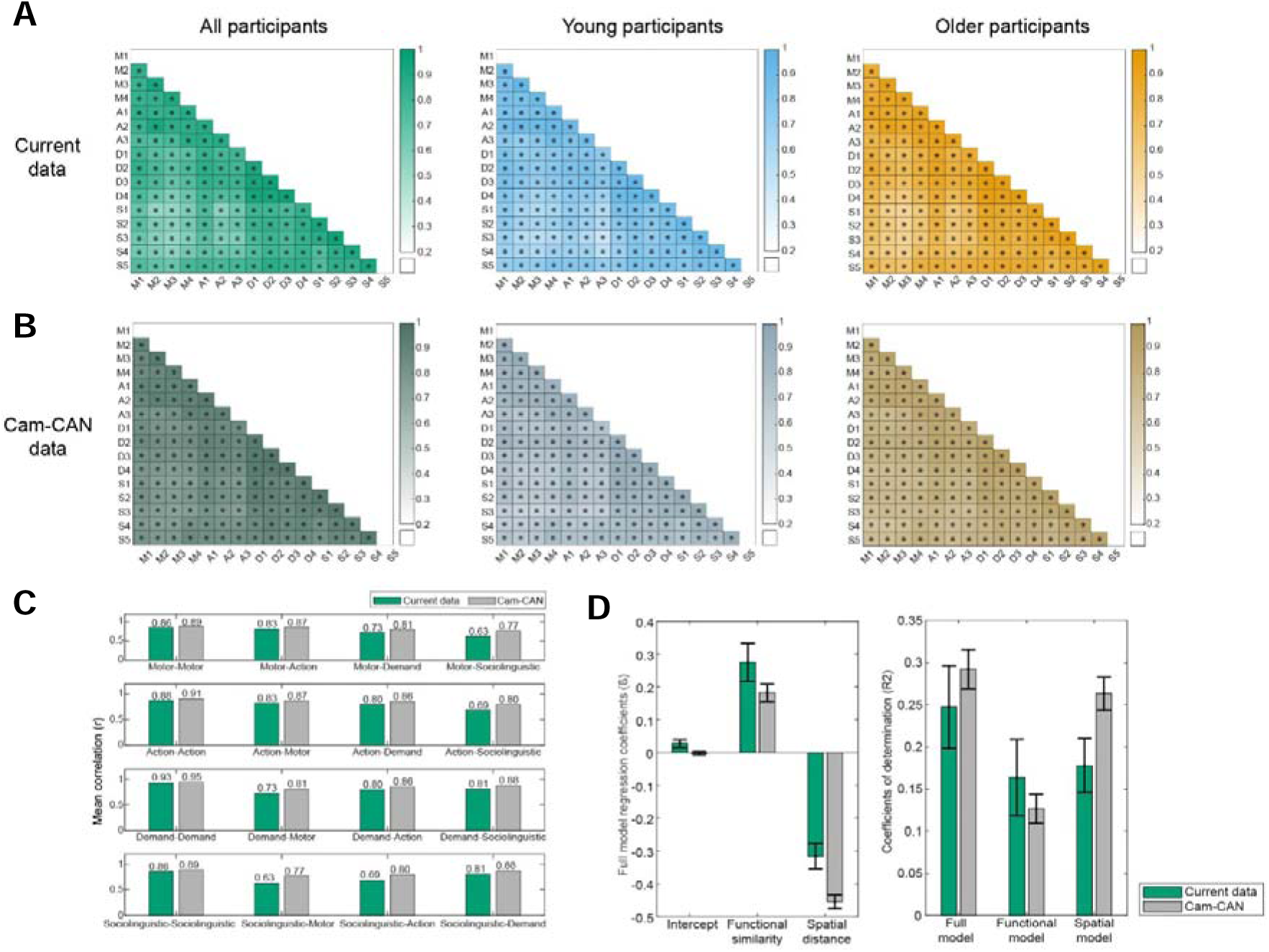
Correlations between functional cerebellar regions. **(A)** Mutual gray matter volume partial correlations between functional cerebellar regions in our dataset, shown for all participants combined (left, green), young adults only (middle, blue), and older adults only (right, orange). Higher correlations appear in more saturated colors, whereas lower correlations appear lighter. Significant correlations are marked with a star. **(B)** Mutual gray matter volume correlations between functional cerebellar regions in the Cam-CAN, shown for all participants combined (left, gray-green), young adults only (middle, gray-blue), and older adults only (right, gray-orange). **(C)** Bar plots show the mean partial correlations within and between each functional domain. Each row begins with the mean within-domain partial correlation, followed by the mean partial correlations between that domain and all other domains. Mean partial correlations based on our own data are shown in green, and those based on the Cam-CAN dataset are shown in gray. **(D)** Left: Bar plots showing the mean beta coefficients from the full model predicting gray matter correlations between regions as a function of spatial distance and functional similarity, for our own dataset (green) and the Cam-CAN dataset (gray). Error bars represent standard deviations. Right: Bar plots showing the mean R^2^ values for the full model, which includes all predictors, and for the reduced models that include each predictor individually, for our own dataset (green) and the Cam-CAN dataset (gray). Error bars represent standard deviations.

In the Cam-CAN dataset, we also observed strong overall correlations (r > 0.73, p < 0.001), and a similar pattern of stronger correlations among regions from the same functional domain (Figure 3B, left). This pattern was consistent across the split young and older adult groups, with no significant differences in correlations between age groups (Figure 3B, middle and right).

In addition, we examined whether regions belonging to the same functional domain showed stronger associations than regions from different domains. This analysis was motivated by the initial observation that mean within-domain correlations appeared higher than between-domain correlations when computed across all participants (Figure 3C, left). To statistically evaluate this, we performed a bootstrap analysis on the partial correlation matrices, comparing within-domain correlations against between-domain correlations for each domain pair. For every functional domain, correlations among regions within the same domain were significantly higher than correlations with regions from other domains, both in the current dataset and in the Cam-CAN dataset (all comparisons: *p* < 0.05, Benjamini– Hochberg corrected).

We further investigated whether higher within-domain correlations compared to between-domain correlations truly reflected functional similarity, or whether this was driven by spatial proximity between regions. We performed a robust multiple linear regression using the partial correlation coefficients as dependent variables, and the functional groupings and Euclidean distance between regions as predictors. The full model included both predictors and explained a substantial proportion of variance in the correlation values (R^2^_full_ = 0.24). Both functional similarity (ß_function_ = 0.28, p_function_ < 0.001) and spatial proximity (ß_spatial_ = −0.32, p_spatial_ < 0.001) significantly predicted the partial correlation coefficients.

We assessed the relative contributions of functional grouping and spatial distance by comparing the full model against the reduced models that included either functional grouping alone or spatial distance alone. For each model, R^2^ values were computed, and their differences were evaluated using a non-parametric bootstrap procedure (5000 iterations), in which the partial correlation matrix was recalculated on resampled participants. This allowed us to test whether one model explained significantly more variance than another. In the current dataset, the full model outperformed both the functional and the spatial model (p < 0.001, Figure 3C, right). The explanatory power of the spatial model (R^2^_spatial_ = 0.18, p < 0.001) was similar to the functional model (R^2^_function_ = 0.16, p < 0.001; spatial vs. functional: *p* = 0.67). Note that an additional analysis showed that spatial distance and functional similarity were negatively correlated (−0.39, p < 0.001), indicating that regions that are more functionally similar also tend to be located closer together. Altogether, these results showed that regions that are functionally similar and spatially close exhibit stronger correlations.

Similar results were found in the Cam-CAN data, in which the full model explained a comparable proportion of variance (R^2^ = 0.29), with all predictors contributing significantly (ß_function_ = 0.18, p < 0.001; ß_spatial_ = −0.45, p < 0.001; Figure 3C, right). The full model performed better dan each of the simple models (both comparisons: p < 0.001, R^2^_function_ = 0.12, R^2^_spatial_ = 0.26). However in contrast to the present results, the spatial model outperformed the functional model for the Cam-CAN dataset (p < 0.001).

### 3.3. Cerebellar-specific task outcomes do not correlate with each other

If there is a link between cerebellar structure and function, we should expect that, given the heterogeneous cerebellar degeneration and the preservation of the topological organization across ages that cerebellar outcomes from tasks linked to the same functional domain would be correlated with each other. To test this, we examined whether the cerebellar outcomes from different tasks were mutually correlated. Across all participants, no significant correlations between any of the tasks were found (Figure 4, left).

**Figure 4.**
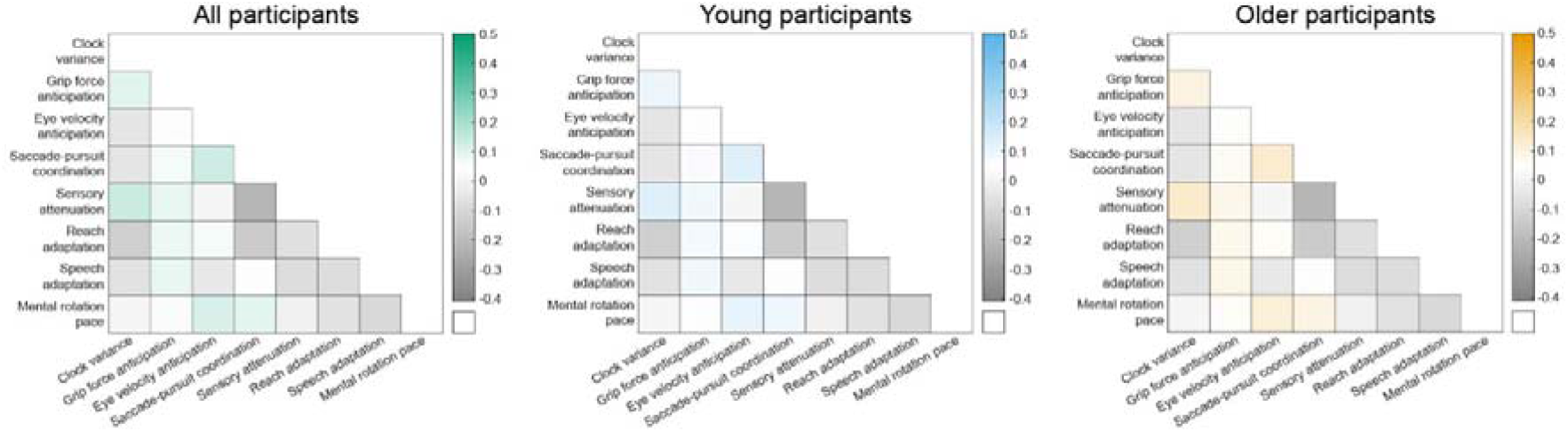
Correlations between cerebellar task performances. Mutual correlations between tasks, shown for all participants combined (left, green), young adults only (middle, blue), and older adults only (right, orange). Strong positive correlations are displayed in the most saturated colors, correlations near zero appear in lighter shades, and negative correlations are shown in gray.

When participants were analyzed separately by age group, initially several associations emerged in the young group before correction. However, none survived the Benjamini-Hochberg correction (Figure 4, middle). In the older adults no significant correlations were observed. Thus, no reliable evidence of associations between cerebellar task performances was found (Figure 4, right).

### 3.4. Regional Gray Matter Volumes do not account for the variability of cerebellar outcomes across participants

We constructed predictive models to test whether gray matter volumes in cerebellar regions could account for individual differences in task performance. The noise ceilings estimated for each task suggested that, in principle, a substantial proportion of the variance in task performance was predictable. Separate models were fit for each task and for each atlas (anatomical and functional). Across all tasks and both atlases, the cross-validated models yielded negative coefficient of determination (R^2^ < 0), indicating that cerebellar gray matter volumes were unable to predict performance (Figure 5A). In other words, the models performed worse than a baseline model that simply predicts each participant’s performance using the mean of each outcome. Additionally, none of the regression coefficients of the predictors, which are the gray matter volumes in the anatomically and functionally defined cerebellar regions, reached statistical significance (Figure 5B). This indicates that gray matter volume of none of the cerebellar regions contributed to predicting task performance.

**Figure 5.**
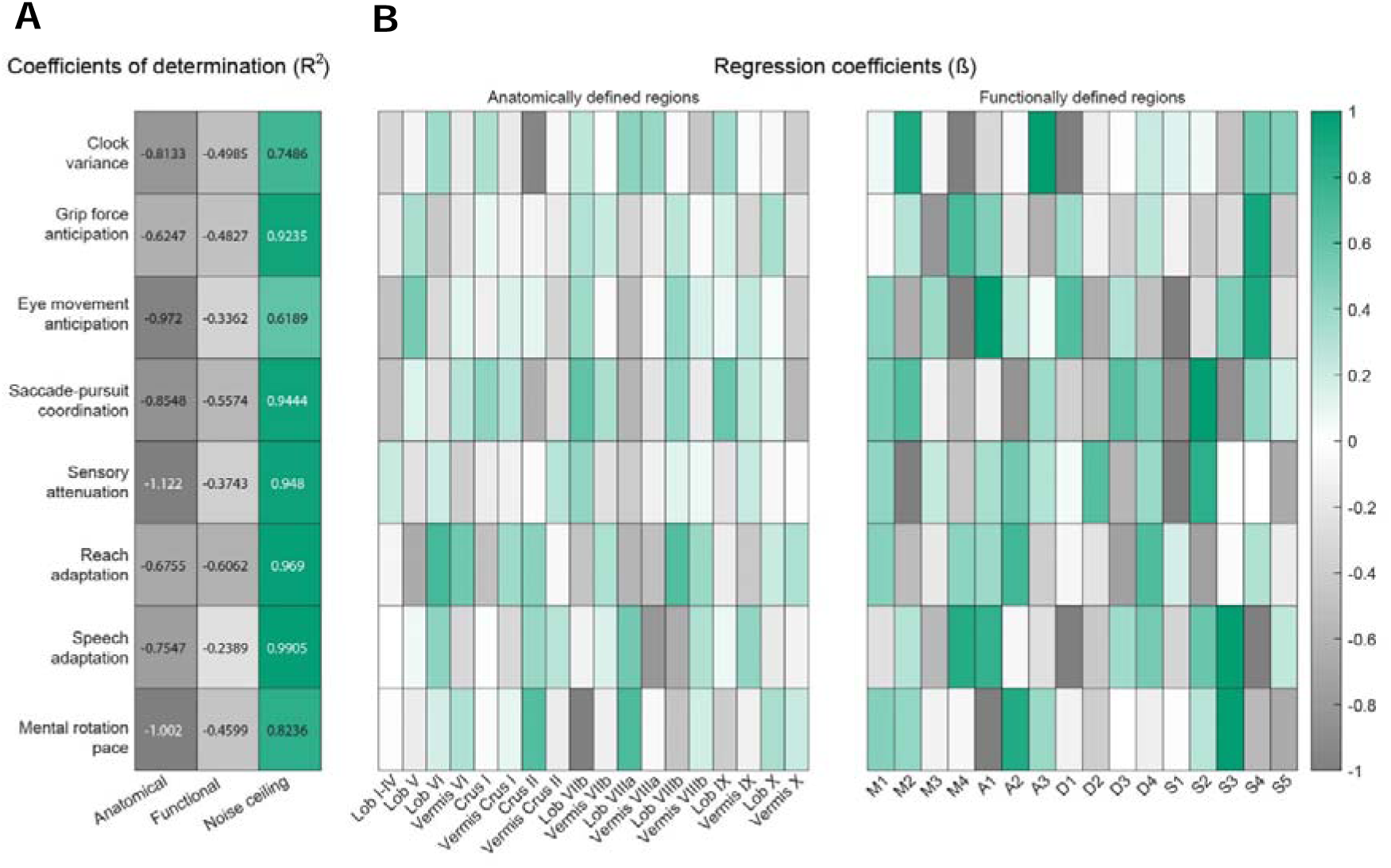
Predictive model results. **(A)** R^2^ values indicate how well the anatomical and functional model fit the task data, relative to the noise ceiling, which represents the maximum explainable variance for each task. Strong positive values are displayed in saturated green, whereas negative values are shown in gray. **(B)** Left: values of the β-coefficients from the anatomical model, in which gray matter volumes of the anatomically defined regions were used as predictors. Right: values of the β-coefficients from the functional model, in which gray matter volumes of the functionally defined regions were used as predictors. None of the predictors did significantly contribute to the model.

Similar patterns were observed when constructing separate predictive models for younger and older adults: the models again yielded negative coefficients of determination, and the regional cerebellar gray matter volumes did not make a significant contribution as predictors (Supplementary Figure 3).

## 4. Discussion

In this cross-sectional study in young and older adults, we assessed the pattern of age-related gray matter degeneration across cerebellar regions and the relation with sensorimotor performance. We found that age-related effects on cerebellar gray matter were diverse both when regions were defined anatomically and when they were defined based on functional parcellations. The structural covariance of cerebellar gray matter was influenced both by spatial proximity but also by the anatomical or functional boundaries of the cerebellum. Yet, gray matter volume in neither anatomically nor functionally defined regions predicted the cerebellar-specific sensorimotor performance in any of the tasks, in none of the populations tested.

### 4.1 Regional gray matter age differences

To characterize patterns of age-related gray matter deterioration in the cerebellum, anatomically defined regions demonstrate larger inter-regional differences in age-related changes in cerebellar structures than functionally defined regions (Figure 1). There was clear variability in the age-related effects on gray matter volumes across anatomically defined regions with posterior regions showing larger age effects than anterior ones. This contrasts with previous studies suggesting that age-related gray matter decline was larger in anterior areas (Bernard and Seidler, 2013; Hulst et al., 2015; Wang et al., 2024), Our results clearly demonstrate across the two datasets that age effects were more pronounced in the posterior lobe than in the anterior one (d’Oleire Uquillas et al., 2026). Interestingly, there is a large overlap in the cerebellar regions with the largest age effects in the study of d’Oleire Uquillas and colleagues and in the present study. In both studies, lobule V and Crus I demonstrated very large age effects. However, lobule VI exhibited very large effects in our study and much smaller ones in the study of d’Oleire Uquilas and colleagues. One can imagine that the technique used to parcellate the different anatomical regions of the cerebellum could account for these differences.

Similarly, the functionally defined regions exhibited different gray matter reductions across the cerebellum in older adults, with the areas of the action domain being the least affected and those of the demand domain the most affected. While these might suggest that some tasks related to the demand domain could be more affected than tasks from the other domains, we caution against such interpretation as no studies have currently provided evidence for such a prediction. Rather, the present study and our previous ones (De Witte et al., 2026a; Matthijs et al., 2026) suggests that cerebellar structure and function are decoupled during the aging trajectory.

Interestingly, the heterogeneous correlations across anatomically defined regions differed from the uniformly high correlations across functionally defined regions (Figure 2A–B versus Figure 3A–B). This suggests that functional parcellations captures regions that are more structurally independent with one another than the anatomically regions are. Overall, these results suggest that there are some differences in age-related changes in gray matter volumes at the anatomical level. The illustration of the effect sizes in Figure 1 suggest indeed the existence of regional differences in vulnerability to aging across the cerebellum.

This interpretation is further supported by the fact that the topological organization of the cerebellum, as assessed by the structural covariance matrices, is highly similar across age groups. If one cerebellar area declined more than the other ones, we would expect the covariance matrices to differ between young and older participants. While the changes in the structural covariance matrix across age have been reported before for cortical regions (Alexander-Bloch et al., 2013; Montembeault et al., 2012; Wu et al., 2012), this study is the first to investigate age-related changes in the topological organization of the cerebellum. In contrast to what was found for the cortex, we did not find any evidence that the topological organization of the cerebellum differed across age groups in our own dataset or in the larger Cam-CAN dataset. Indeed, none of the correlations between the gray matter volumes of different cerebellar regions differed significantly.

Interestingly, our modeling approach revealed that the topological organization of the cerebellum was partially determined by the functional domains and by the lobular boundaries for both young and older participants. The functional parcellation revealed a significant effect of functional domain on the topological organization as gray matter correlations were higher between regions belonging to the same functional domain (action, motor, demand, or sociolinguistic) compared with correlations between regions from different domains (Figure 3C). Moreover, in the model, functional similarity derived from continuous task-based activity patterns reported by Nettekoven et al. (2024) also substantially explained inter-regional gray matter correlations. This suggests that the functional parcellation is not only interesting to investigate functional patterns but also to look at structural changes in the cerebellum.

As in the analyses of the topological organization of the cortex, structural covariance was highly influenced by spatial proximity of the correlated regions. In both anatomically and functionally defined parcellations, spatial proximity between regions emerged as a robust predictor of the gray matter correlations observed between them. Notably, within the anatomical parcellation, spatial proximity explained inter-regional gray matter correlations more effectively than either lobular or medial–lateral divisions of the cerebellum. In our dataset, the lobular division (anterior vs. posterior) could not account for the patterns observed in the correlation matrices. In addition, the anterior-posterior model always explained a smaller portion of variance than the full model, which reinforces the fact that this division might not be as relevant for age-related effects than previously claimed. In contrast, the medial-lateral division (vermis vs. hemisphere) could account for the patterns of the structural covariance matrix for both datasets. This indicates that the topological organization of the cerebellum transcends strict anatomical boundaries, such as the primary fissure, and is better explained by the spatial proximity of the different anatomically regions. Consequently, it underscores a fundamental drawback of relying on coarse predefined regional parcellations, such as vermis versus hemispheres or anterior versus posterior regions, which may be too general to serve as effective predictors, as they encompass large and functionally heterogeneous areas, potentially obscuring more localized patterns of age-related change.

One limitation of this analysis is the strong correlation between spatial distance and functional similarity, which raises the possibility that these predictors may serve as different proxies for a shared underlying organizational process. The relative weight of the spatial proximity and the functional domains varied across both datasets. Results from the larger Cam-CAN dataset showed that spatial distance could have a larger weight than the functional similarity as a determinant of structural covariance. Nevertheless, in both datasets, combining spatial distance and functional similarity substantially improved predictive performance than each of these predictors alone. This indicates that each predictor captures partly distinct aspects of the underlying signal, demonstrating a unique contribution of functional similarity in addition to the dominant effect of spatial proximity.

Interestingly, the variance accounted for by the different models was larger for the functional parcellation (R^2^ ~0.3) than for the anatomical parcellation (R^2^ ~0.1). Spatial distance accounted for more variance in structural covariance among functional regions (R^2^~0.2) than among anatomically regions (R^2^~0.1). It is unclear what this difference in explanatory power means. The lower variance accounted for by the anatomical parcellation suggests that other yet unknown factors such as microstructural boundaries, or simply differences introduced by the parcellation itself, play a larger role in determining the correlation structure. However, it is important to note that the explanatory power of all predictive models remained modest (R^2^ < 0.3; Ben-Shachar et al., 2020), indicating that most of the variance in inter-regional gray matter correlations is not accounted for by our spatial and functional determinants thus remains unexplained by this study.

### 4.2 Absence of structure-behavior relation

The fact that age-related effects on cerebellar gray matter were quite heterogeneous across functionally and anatomically defined regions, but that the gray matter volumes of these different functional regions were highly correlated suggests that we might predict changes in function based on gray matter changes. This could account for the fact that cognitive function declines faster than motor functions.

Given the maintenance of the topological organization of the cerebellum with age and if cerebellar structure determines its function, we should expect that cerebellar outcomes from our tasks would be correlated with each other, especially in older participants. Yet the outcomes of our eight task performances were not correlated with each other (Figure 4). This finding supports the notion that distinct cerebellar mechanisms underlie different tasks, rather than a single latent cerebellar factor across tasks (Löwgren et al., 2020). In our previous study, we found that older adults performed as well as younger adults on these tasks, despite exhibiting reduced cerebellar volumes (De Witte et al., 2026a; Matthijs et al., 2026, 2025). This suggests that the deterioration of the cerebellar structure (e.g. total cerebellar volume) cannot account for the maintenance of cerebellar function.

To test the uncoupling idea further, we examined, in this study, the extent to which the inter-participant variability in cerebellar outcomes from these eight sensorimotor tasks could be explained by gray matter volumes of different cerebellar regions, using either anatomical and functional parcellations. Our results showed that performance on none of the eight sensorimotor tasks could be accounted for by gray matter volume of anatomical or functional cerebellar volumes (Figure 5). In other words, inter-individual differences in cerebellar-dependent sensorimotor behavior are not driven by structural variability within specific cerebellar regions. This could either mean that none of the “cerebellar” outcomes actually depend on the cerebellum (a possibility that is extensively discussed in De Witte et al., 2026a) or that cerebellar structure and function are not directly related to each other.

The absence of a structure–behavior relationships was particularly unexpected for the functionally defined parcellations. These parcellations delineate cerebellar regions based on task-evoked functional activity patterns (Nettekoven et al., 2024), and therefore we hypothesized a closer correspondence between regional gray matter volume and task performance associated with those same functions. A plausible explanation for the lack of such a relationship is that the functional parcellation atlas was derived primarily from functional activity profiles rather than from structural features such as gray matter. As a result, these functionally defined boundaries may be optimally suited for capturing task-related patterns of neuronal activation, but not for identifying structural variability that would meaningfully predict behavioral differences.

A particularly compelling aspect of our results is the consistency of the null effect across all eight sensorimotor tasks. The absence of any detectable relationship between regional gray matter volume and task performance was not limited to a subset of tasks but was observed uniformly, regardless of whether regions were defined anatomically or functionally. This convergence across multiple independent measures strengthens the conclusion that regional gray matter metrics do not explain variance in cerebellar-dependent sensorimotor behavior. Moreover, the negative R^2^ values indicate that this pattern is unlikely to reflect insufficient statistical power, but rather a genuine absence of structure-function relationship, as low statistical power would typically yield R^2^ values fluctuating around zero. Instead, consistently negative values point to a genuine absence of predictive signal in the data. Combined with high noise ceilings for each task, suggesting that a considerable portion of behavioral variance could, in principle, be accounted for, these findings provide robust evidence that structural volume fails to capture the functional complexity underlying cerebellar contributions to sensorimotor control.

This finding is in contrast with previous results showing that working memory, balance and choice reaction time performances could be predicted by regional cerebellar gray matter volumes (Bernard and Seidler, 2013). One explanation is that we did not separate left and right cerebellar hemispheres in our analysis. While age-related effects are generally similar across hemispheres, they differ in absolute volume, which could, in principle, mask hemisphere-specific contributions; however, given the overall correlation across cerebellar regions, any such influence is likely to be modest (Bernard and Seidler, 2013). On the other hand, these tasks may involve greater cognitive demands compared to our primarily sensorimotor measures, except for the mental rotation task. Supporting this idea, Nadkarni et al. (2014) demonstrated that the relationship between motor performance and cerebellar volume disappears when controlling cognitive factors such as information processing speed. This may suggest that cognitive functions depend more strongly on cerebellar structure than motor functions do; however, direct empirical evidence for this interpretation remains limited and our data argues against this possibility as neither motor nor cognitive cerebellar outcomes could be explained by the structure of the cerebellum.

Another explanation for the absence of structure–function associations in our results is the reliance on predefined regions of interest based on morphological features (anatomically defined regions) or activity-derived parcellations (functionally defined regions). These approaches assume that the boundaries of cerebellar organization align neatly with either anatomical landmarks or functional activation patterns, which may not fully capture the distributed and fine-grained nature of cerebellar contributions to behavior in the tasks we used. It remains plausible that a structure–behavior relationship could emerge when gray matter volume is assessed at a voxel-wise level, allowing for the detection of localized clusters that do not conform to predefined regional boundaries. Future studies should confirm this by employing voxel-based morphometry or multivariate pattern analysis, which may uncover associations that were masked by region-level averaging in this study.

Together, these findings may also be understood in the context of the concept of cerebellar reserve (Arleo et al., 2023; Bordignon et al., 2021; Gelfo and Petrosini, 2022; Mitoma et al., 2020). This framework proposes that the cerebellum possesses a robust capacity to compensate for structural loss through functional reorganization and distributed network engagement (Coello et al., 2026). The previously reported absence of age-related differences in task performance between young adults, older adults between 55 and 70 years old and even older old adults above 80 years old, despite reductions in cerebellar volume in older and older adults compared to younger adults (De Witte et al., 2026b), combined with the present lack of structure–function associations between cerebellar regions and task performance, suggests that regional gray matter volume may not be the primary determinant of behavioral variability. One possibility is that, although cerebellar gray matter volume is reduced in our sample, it may remain sufficiently large to support efficient functional performance, resulting in a ceiling effect beyond which variation in volume no longer predicts behavioral differences. Alternatively, sensorimotor performance may be maintained through functional redundancy, adaptive network recruitment, or microstructural properties that remain preserved even in the context of overall age-related volume reductions (Cabeza et al., 2018; Sarah G. Donofrio et al., 2025; Morcom and Johnson, 2015). Together, these considerations align with the notion that cerebellar reserve enables stable behavioral performance despite substantial structural change.

### 4.3 Conclusion

Age-related effects on the cerebellum appear to be variable across anatomical and functionally defined regions. However, the topological organization of the cerebellum remains the same across age groups and seems to be driven both by spatial proximity, and by medial-lateral division or functional similarity. These findings indicate that, even in the presence of age-related degeneration, the cerebellum’s structural organization remains highly stable. Finally, gray matter volume of neither anatomically nor functionally defined cerebellar regions appear to account for inter-individual variability in cerebellar task outcomes, which suggests that the variability in cerebellar gray matter has poor explanatory power for the variability in cerebellar behavior. Taken together, these findings are consistent with the concept of cerebellar reserve, suggesting that cerebellar function can be maintained across aging despite degeneration of regional gray matter structure.

## 5. Supplementary methods

### 5.1 Cam-CAN data: linear relationship between age and structure

We analyzed the full Cam-CAN cohort to assess the linear relationship between gray matter volume and age. For this, we used MATLAB’s *polyfit* function with a polynomial degree of 1, which fits a first-degree polynomial to the data using a least-squares approach. This yields a linear regression line modeling changes in volume with age. The analysis was conducted separately for each functional cerebellar region as well as each anatomical region.

## 6. Supplementary results

### 6.1 Euclidean distances between regions

**Supplementary Table 1.**
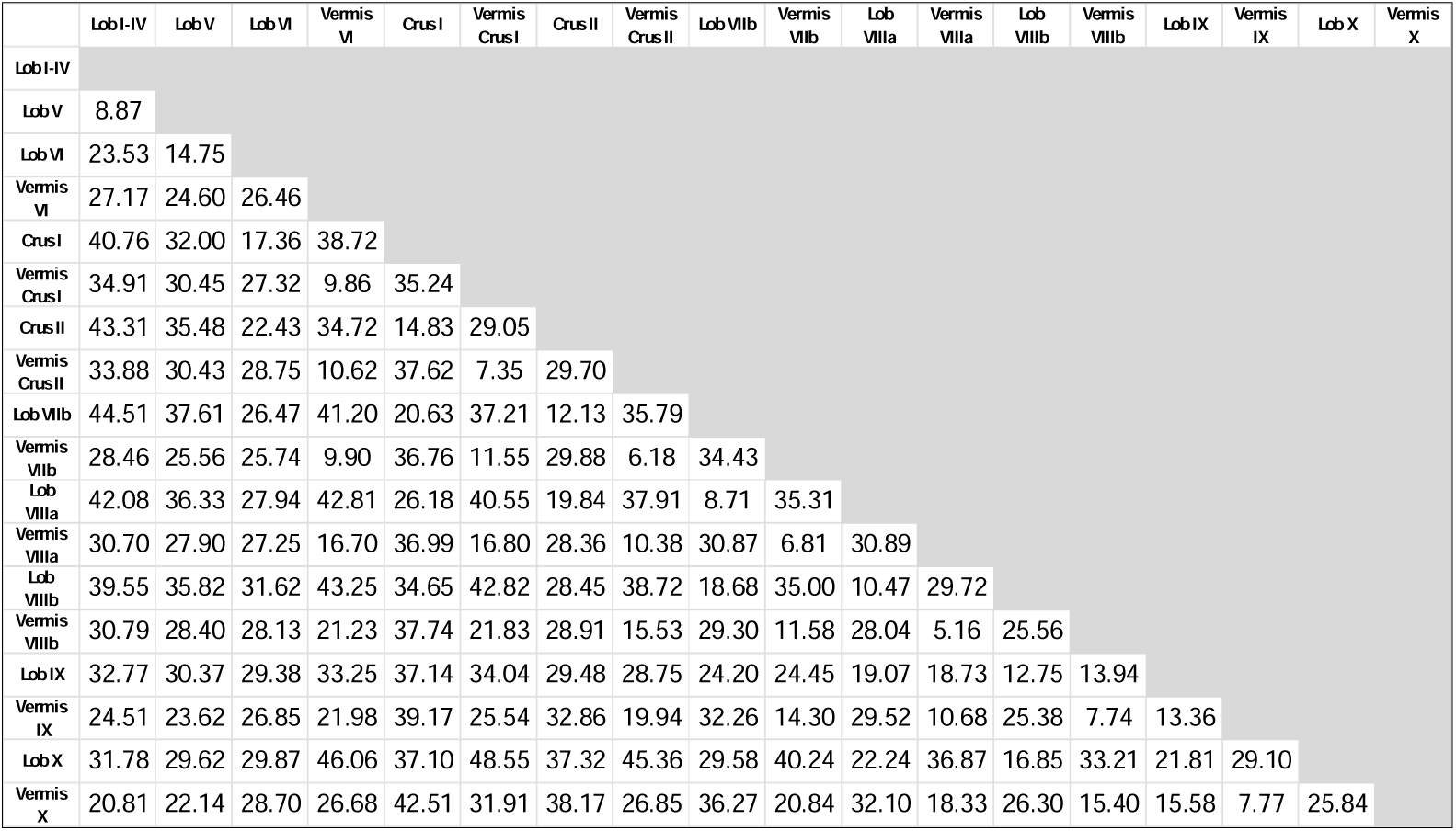
Euclidean distances between anatomically defined regions.

**Supplementary Table 2.** Euclidean distances between functionally defined regions.

|  | M1 | M2 | M3 | M4 | A1 | A2 | A3 | D1 | D2 | D3 | D4 | S1 | S2 | S3 | S4 | S5 |
| --- | --- | --- | --- | --- | --- | --- | --- | --- | --- | --- | --- | --- | --- | --- | --- | --- |
| M1 |  |  |  |  |  |  |  |  |  |  |  |  |  |  |  |  |
| M2 | 15.34 |  |  |  |  |  |  |  |  |  |  |  |  |  |  |  |
| M3 | 15.03 | 8.24 |  |  |  |  |  |  |  |  |  |  |  |  |  |  |
| M4 | 23.80 | 15.76 | 11.33 |  |  |  |  |  |  |  |  |  |  |  |  |  |
| A1 | 15.80 | 12.70 | 14.35 | 26.61 |  |  |  |  |  |  |  |  |  |  |  |  |
| A2 | 18.70 | 9.72 | 7.54 | 9.85 | 19.16 |  |  |  |  |  |  |  |  |  |  |  |
| A3 | 23.07 | 11.14 | 9.30 | 10.34 | 17.50 | 7.93 |  |  |  |  |  |  |  |  |  |  |
| D1 | 15.95 | 13.90 | 14.22 | 22.82 | 10.72 | 14.85 | 17.27 |  |  |  |  |  |  |  |  |  |
| D2 | 23.73 | 18.71 | 16.60 | 17.75 | 23.52 | 14.63 | 16.27 | 16.30 |  |  |  |  |  |  |  |  |
| D3 | 19.56 | 10.24 | 13.79 | 18.61 | 14.22 | 10.78 | 12.61 | 10.34 | 14.84 |  |  |  |  |  |  |  |
| D4 | 24.20 | 19.61 | 17.79 | 22.58 | 19.77 | 15.52 | 18.20 | 11.57 | 7.36 | 10.76 |  |  |  |  |  |  |
| S1 | 12.21 | 16.30 | 18.78 | 26.44 | 12.76 | 20.63 | 22.86 | 11.24 | 21.40 | 15.74 | 19.54 |  |  |  |  |  |
| S2 | 14.27 | 19.81 | 19.68 | 26.13 | 16.14 | 21.19 | 24.66 | 12.14 | 18.99 | 16.92 | 14.86 | 8.50 |  |  |  |  |
| S3 | 18.00 | 21.45 | 20.41 | 25.61 | 20.29 | 21.82 | 25.26 | 14.63 | 14.75 | 18.83 | 14.01 | 13.52 | 8.78 |  |  |  |
| S4 | 28.12 | 22.78 | 20.68 | 21.54 | 26.02 | 18.13 | 22.64 | 19.69 | 8.27 | 15.86 | 10.24 | 24.31 | 20.02 | 15.35 |  |  |
| S5 | 32.86 | 21.37 | 19.91 | 12.23 | 31.09 | 13.80 | 15.47 | 24.86 | 16.60 | 18.77 | 19.98 | 30.72 | 29.32 | 30.80 | 19.26 |  |

### 6.2 Regression between regional gray matter and age in Cam-CAN data

In addition to the cross-sectional comparison between young and older Cam-CAN participants, we performed a regression analysis between age and gray matter volume across the full Cam-CAN cohort to visually assess whether the cross-sectional pattern reflects the overall age-related trend. For the anatomical regions, Cam-CAN data revealed a negative correlation between gray matter and age across all regions (Supplementary Figure 1). Overall, these results were consistent with the cross-sectional results from this sample. The steepest declines were observed in lobule VI, Crus I, and Crus II, regions that also showed strong effect sizes in our data, while the shallowest declines occurred in vermis X and Vermis Crus II, which corresponded to regions with weaker effect sizes in our sample. Notably, Cam-CAN data indicated only a weak slope in Vermis Crus I, despite this region showing a strong age-related effect in our sample. This was also shown in the weaker Cam-CAN effect size compared to our own for this region (**Error! Reference source not found.** A) and is attributable to the very small size of this region, which makes parcellation results less reliable.

The functional regions also showed negative correlations across all areas, consistent with the results from our dataset. The regression slopes across regions exhibited highly similar patterns (Supplementary Figure 2).

**Supplementary Figure 1.**
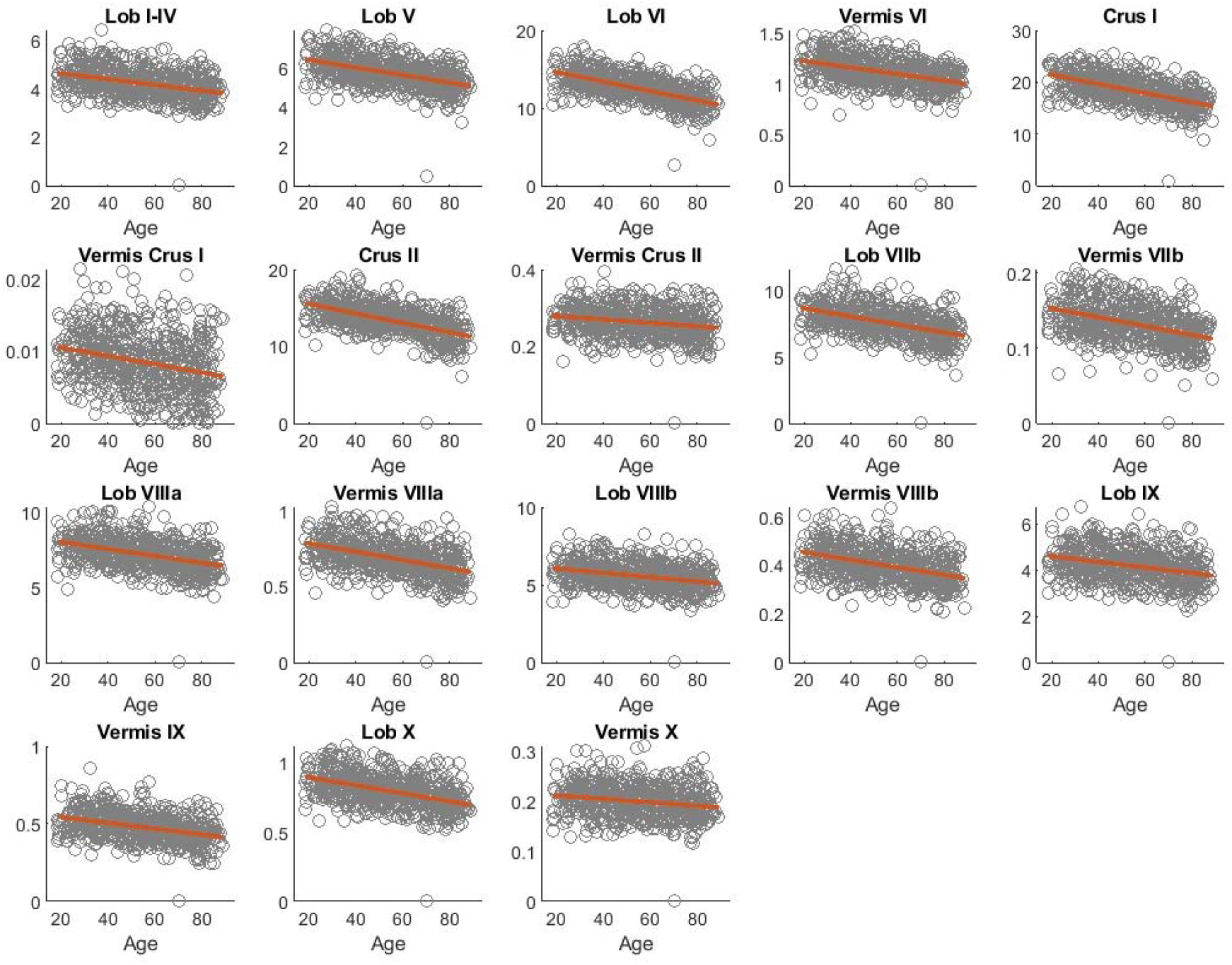
Regression results between age and gray matter volume in each anatomically defined region of the cerebellum in the Cam-CAN dataset. The orange line represents the regression slope, and the gray, unfilled scatter points show the data from individual participants.

**Supplementary Figure 2.**
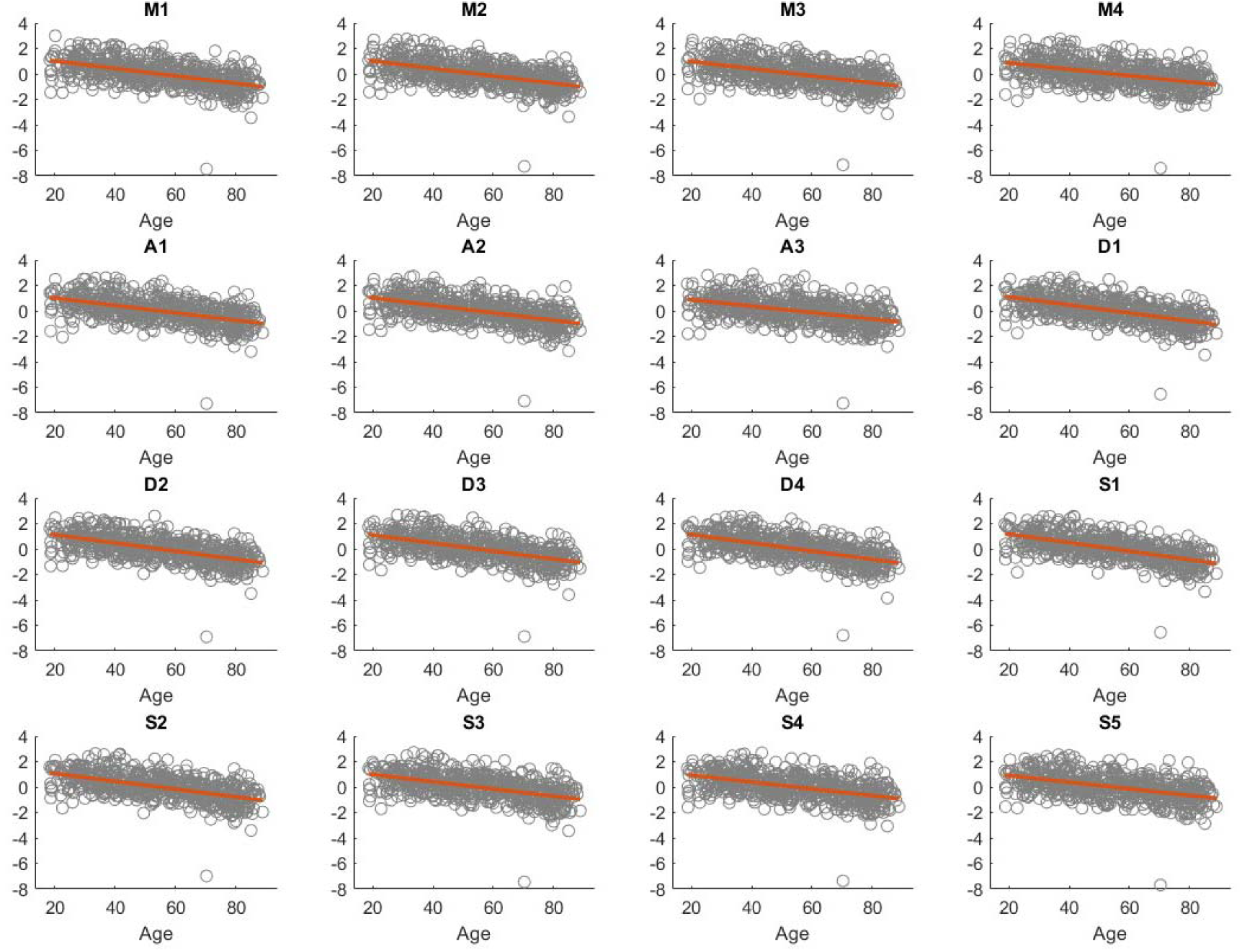
Regression results between age and gray matter volume in each functionally defined region of the cerebellum in the Cam-CAN dataset. The orange line represents the regression slope, and the gray, unfilled scatter points show the data from individual participants.

### 6.3 Predictive models per age group

The models predicting task performance from regional gray matter volume for all participants combined showed limited predictive power (**Error! Reference source not found.**). We additionally created separate models for the young and older groups to assess whether similar results were obtained within each age group. In the young participants, none of the models were able to predict task performance on any of the tasks (R^2^ < 0; Supplementary Figure 3A). Additionally, none of the regression coefficients associated with the individual anatomical and functional cerebellar regions reached statistical significance (Supplementary Figure 3B). This was also the case for the older participants (Supplementary Figure 3C and D).

**Supplementary Figure 3.**
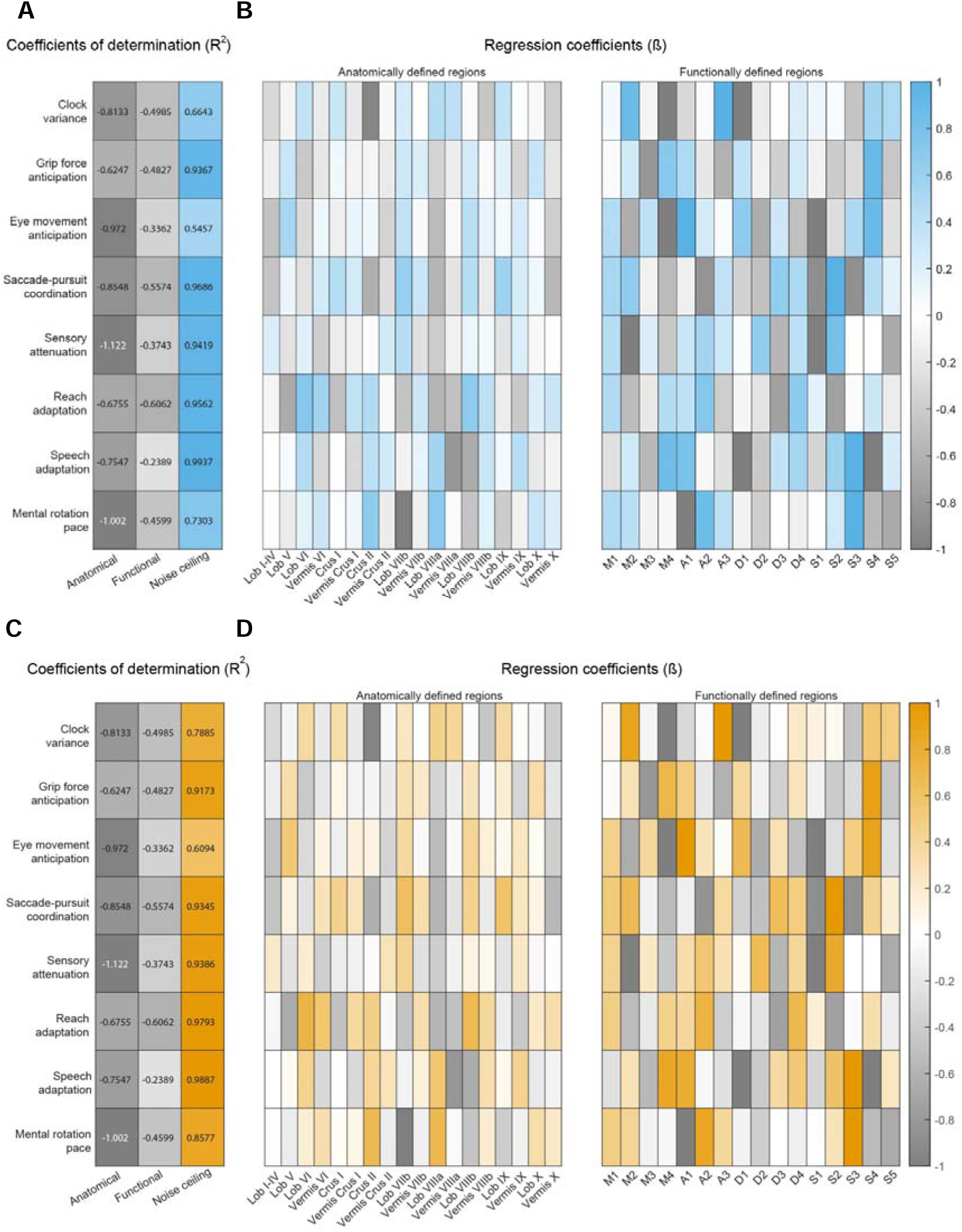
Predictive model results for young and older participants. **(A)** R^2^ values indicate how well the anatomical and functional model fit the task data for the young participants, relative to the noise ceiling, which represents the maximum explainable variance for each task. Strong positive values are displayed in saturated color, whereas negative values are shown in gray. **(B)** Left: values of the β-coefficients from the anatomical model for young participants, in which gray matter volumes of the anatomical regions were used as predictors. Right: values of the β-coefficients from the functional model for young participants, in which gray matter volumes of the functional regions were used as predictors. **(C)** R^2^ values indicate how well the anatomical and functional model fit the task data for the older participants, relative to the noise ceiling. **(D)** Left: values of the β-coefficients from the anatomical model for older participants. Right: values of the β-coefficients from the functional model for older participants.

## Acknowledgment

This work was supported by the Fonds Wetenschappelijk Onderzoek (FWO) (G095121N awarded to JJO). C.R.N. was supported by a Wellcome Trust Early Career Award (306553/Z/23/Z).

The CamCAN dataset was provided by the Cambridge Centre for Ageing and Neuroscience (CamCAN). CamCAN funding was provided by the UK Biotechnology and Biological Sciences Research Council (grant number BB/H008217/1), together with support from the UK Medical Research Council and University of Cambridge, UK.

## Disclosure statement

None of the authors reported a conflict of interest.

## Notes

### Competing Interest Statement

The authors have declared no competing interest.

### Summary of Updates

In this version, we updated the results as we discovered a mistake in the code used to analyse the data. While, in a previous version, we mentioned that age-related difference in cerebellar gray matter was uniform, we now say that this difference varies across cerebellar regions.

